# Global characterization of *Dictyostelium discoideum* gene expression changes under hypoxic conditions

**DOI:** 10.1101/2025.04.02.646759

**Authors:** Julie Hesnard, Elisabet Gas-Pascual, Hanke van der Wel, Olivier Cochet-Escartin, Stephane Joly, Jean-Paul Rieu, Christopher M West, Christophe Anjard

## Abstract

Aerobic eukaryotes utilize O_2_ to oxidize metabolites and generate ATP. The protist kingdom lacks the Hypoxia Inducible Factor-dependent transcriptional response network to accommodate low O_2_ though possesses a HIF prolyl hydroxylase that initially mediates the response. To address the scale and scope of hypoxic responses in protists, we characterized transcriptomic and proteomic changes when the social amoeba *Dictyostelium* is subjected to low (1%) O_2_ under nutritive conditions over 24 h followed by reoxygenation. Remarkably, 32% of the transcripts quantified were differentially expressed during hypoxia, with greatest changes associated with early (1 h) and late phases (24 h). Protein changes were modestly correlated with and generally lagged behind transcriptional changes. Correlated changes were observed for transcripts and proteins associated with various metabolic, anabolic, and catabolic pathways, as well as chromosome organization, cell cycling, vesicular trafficking, and signaling. Analysis of 4 marker genes showed extremely rapid responses that were graded over a range of O_2_ levels, with differential responses to inhibitors affecting protein synthesis and mitochondria. Overall, the amoebal response to a low but non-toxic O_2_-level resulted in massive remodeling of the transcriptome and proteome.

## Introduction

Most eukaryotes on Earth’s surface are aerobic, using atmospheric oxygen (O_2_) to generate the energy needed for their metabolism (1). However, the concentration of oxygen dissolved in environmental, organismic and cellular waters varies dramatically from essentially zero to 21% or ∼200-250 µM (2,3). Organisms and the cells within them have evolved their O_2_- dependent enzymes and O_2_-binding proteins to function within the variations that they normally encounter. The ability of cells to sense in addition utilize O_2_ for energy production can provide information regarding their location within O_2_ concentration gradients, and/or to adapt to the change. The responses may lead to migration to locations with more favorable concentrations (aerotaxis), differentiation, metabolic reprogramming, or more global stress responses. Changes in gene expression are particularly well-studied in metazoans using low O_2_ environments (4–6). It was shown that under normoxic conditions, prolyl hydroxylases (PHDs) use O_2_ to hydroxylate the transcription co-factor HIF-1α (7). This modification allows von Hippel-Lindau protein binding, poly-ubiquitination and degradation of the transcription co-factor (8–10). Low O_2_ inhibits hydroxylation, allowing HIF-1α to dimerize with HIF-1β, enter the nucleus and regulate the transcription of thousands of genes (11,12). Many target genes are up-regulated, being involved in angiogenesis, erythropoietin production, glycolysis enhancement and mitochondrial oxygen consumption reduction (13–15). HIF-1 also induces transcriptional regulators such as O_2_-dependent *Jumonji-C* lysine demethylases involved in epigenetic hypoxic regulation (16,17). Additionally, HIF-1 is often overexpressed in cancer cells and contributes to drug resistance, underscoring the importance of understanding hypoxic regulation for effective therapeutic treatments (18–20). Recent studies revealed that some mammalian hypoxic responses are also HIF-independent, highlighting the complexity of this regulation. Transcription factors like Myc and SREBP2, which control respectively glutamine metabolism and cholesterol synthesis during hypoxia, are part of these alternative pathways (21,22). The mechanisms involved and the target genes in these regulations are not yet fully understood. Moreover, most studies are performed on whole organisms or tissues, but transcriptional responses to hypoxic conditions at the unicellular level is still poorly understood.

The social amoeba *Dictyostelium discoideum* is an obligate aerobic organism that lives in the soil, feeding on bacteria (23). When starvation occurs, individual amoebae can aggregate to form multicellular slugs that migrate to the soil surface to culminate into terminally differentiated fruiting bodies with dormant spores perched atop cellular stalks, allowing efficient spore dispersal (24). O_2_ is an important signal for *Dictyostelium*, by influencing single cell aerotaxis (25–27), slug migration (28), slug polarity (29), culmination (30), and terminal spore differentiation (31). *Dictyostelium* lacks the HIF-1 transcription factor but has an O_2_-sensing system based on PhyA (32), likely the protist ortholog of PHD2 (33). The role of this O_2_ sensor was mainly studied during multicellular development, revealing that PhyA controls the O_2_ threshold for culmination leading to fruiting body formation, ensuring that fruiting bodies differentiate at the O_2_-rich surface of the soil to facilitate future dispersal of spores (32,34). At higher O_2_, PhyA hydroxylates a specific proline residue on Skp1, an adaptor subunit of the Skp1–Cul1–F-box (SCF) class of E3 ubiquitin ligases. This modification leads to Skp1 glycosylation, resulting in selective representation of certain F-box proteins (FBPs) with Skp1 and in SCF complexes (35). This mechanism modulates the abundance of affected FBPs, which result from autoubiquitination in SCF complexes. Indeed, the absence of PhyA is partially rescued by treatment of cells with proteasome inhibitors. It is predicted that low O_2_ modulate tagging of target proteins with polyubiquitin chains that mark them for recognition and degradation by the 26S-proteasome. Below 10% O_2_, PhyA activity is reduced, allowing differential remodeling of the proteome. However, the target proteins and their roles in O_2_ adaptation remain unknown. This mechanism appears to be widely conserved among protists, including the pathogens *Toxoplasma gondii* and *Pythium ultimum* (36–38). Independently, it was described in vegetative cells that hypoxia induces a switch from cytochrome C oxidase optional subunit CgxE to CxgS, resulting in a threefold increase in affinity for O_2_ (39). It was also shown that the two flavohemoglobins, FhbA and FhbB, are induced when cells are incubated at 0.4% O_2_, presumably to detoxify nitric oxide produced during hypoxia (40,41). However, the global regulation and the biological processes involved in the hypoxic response of *Dictyostelium* and other protists remain largely unexplored.

The aim of this study is to identify new mediators of O_2_ responses, by exploring the transcriptional and proteomic responses of vegetative stage (proliferating) *Dictyostelium discoideum* amoebae to 1% (12 µM) O_2_. Given O_2_ levels expected in its soil environment, this level will be referred to as hypoxic, in contrast to normoxic which will refer to the 21% (∼240 µM) O_2_ experienced at the soil surface. Transcriptomic and proteomic assays were performed over a time course of 24 h followed by 5 h of reoxygenation. Bioinformatic analyses of the hypoxic responses identified several functional clusters, and revealed an acute expression shift occurring within 1 hour transcriptionally or 5 h proteomically, and a slower, chronic adaptation occurring at 24 h that mostly remains after reoxygenation. RT-qPCRs of select hypoxia marker genes mapped temporal and concentration dependence on O_2_ levels. Furthermore, effects of inhibitors of mitochondrial respiration, *de novo* protein synthesis and the broader class of non-heme dioxygenases were evaluated.

## Materials and methods

### Cell culture and strains

*Dictyostelium discoideum* AX3 cells were provided by the National BioResource Project (NBRP Nenkin, Japan) and cultured axenically with HL5 medium (HLF2; Formedium, UK) containing 100 mM glucose and 1/1000 penicillin-streptomycin (P/S) (15140-122; Gibco, USA). Cells were grown at 22 °C under agitation at 180 rpm.

### Experimental setup used for transcriptomic and proteomic time course

Exponentially growing cells were diluted to ∼2.5 × 10^5^ cells/ml in flasks containing 150 ml (proteomic) or 30 ml (transcriptomic) degassed HL5 medium (Fig. S1). Shaking flasks were continuously and serially purged with a humidified atmosphere of 1% O_2_ in N_2_ for 24 h, and then reoxygenated with 20% O_2_ for 5 h. A flask was removed to collect cells at 1 h, 5 h, and 24 h under hypoxia, and at 1 h and 5 h during reoxygenation. The standard t0 was collected from the culture tube. After cell collection, cell density was evaluated using a hemocytometer at 5 h, 24 h, and 5 h reoxygenation. For proteomics analyses, a parallel set of samples maintained at 21% O_2_ was collected.

2x10^6^ or 10^7^ cells were collected for the transcriptomic or proteomic analyses, respectively, by rapid centrifugation at 1200 × *g* for 1 min at 4°C, resuspension in ice-cold degassed 33 mM NaH_2_PO_4_, 10.6 mM Na_2_HPO_4_, 20 mM KCl, 6 mM MgSO_4_, pH 5.8, and repeated centrifugation. After supernatant aspiration, cell pellets were snap-frozen in liquid N_2_ and stored at -80°C. Each experiment was performed in triplicate (n=3).

### Library preparation and RNA sequencing (Massive Analyses of cDNA Ends from GenXpro)

Total RNAs were extracted (RNeasy Plus Mini Kit, 74134, Qiagen, Germany) and quantified. 100 ng were used for cDNAs synthesis using oligo(dT) primers following the manufacturer’s instructions (Rapid MACE-seq kit, GenXpro GmbH, Germany). The library was amplified by PCR, followed by a SPRI bead clean-up (M1378-00 Mag-Bind TotalPure NGS Omega Bio- Tek, USA). Sequencing and bioinformatic analyses were performed by GenxPro. Samples were sequenced using the Illumina platform NextSeq 500 (5 M reads 1x75 bps). The reads were mapped to Ensembl *Dictyostelium discoideum* 2_7 as a reference genome. Data were analyzed using DESeq2 (version 1.38, (42)) and HTSeq (version 2.0.2) was used to quantify the transcripts. The mean transcripts per million (TPM) from the triplicates was used to calculate the log2 fold changes (log2FC) for each time point, using t0 (21% O_2_) as the reference. The p-values were calculated using the Wald test and corrected with the Benjamini-Hochberg method to obtain the false discovery rate (FDR).

### Turn over calculation

For all detected transcripts, TPM values, provided in Table S1A, were used to calculate turnovers. For each gene, TPM values between two consecutive time points were subtracted to quantify the extent of degradation or synthesis. Then the absolute values of all TPM differences were summed. Finally, the sum was normalized to a ratio of 1 million and converted into a turnover percentage. For each turnover calculation, the mean value of the replicates and the standard deviation were calculated.

### Differential transcript expression analyses

To analyze differentially expressed transcripts, sequences assigned as non-coding RNAs, pseudogenes, mobile elements and tRNAs were removed manually, and the resulting coding RNAs are summarized in Table S1B. Pearson correlation was used to assess continuity of total coding RNAs between time points. An ad hoc code was applied to remove genes with at least one missing expression during the time course. The code was used to categorize genes into up-regulated (log2FC≥1) and down-regulated (log2FC≤-1) groups. Only genes with an FDR<0.05 were considered. Within each regulation group, differentially expressed transcripts were classified based on their GO annotations from DAVID, using dictyBase_IDs (43,44). Only transcripts with significant direct GO term classification (p-value<0.05) and fold enrichment (FDR<0.05) values were included in the analysis. Interactions between encoded proteins from these enriched transcripts were visualized using STRING (version 12.0). Additional pathways were analyzed manually as indicated.

### Comparison with published transcriptomic data

Coding RNA from Table S1B were compared to the transcriptomic data from Lamrabet et al. recent study (45). Total identified RNA data were extracted from the supplement (https://www.frontiersin.org/journals/microbiology/articles/10.3389/fmicb.2020.00410/full) and converted in log2FC. Differentially expressed transcripts (log2FC≥1 or log2FC≤-1; FDR<0.05) after 1 h of hypoxia compared to t0, and differentially expressed transcript (log2FC≥1 or log2FC≤-1; p.value adj<0.05) compared to cells cultured without bacteria, were analyzed.

### Sample preparation for proteomic analyses

Whole cell protein lysates were obtained by SDS-lysis followed by reduction, alkylation, and trypsin digestion according to the S-Trap micro column manufacturer’s recommended digestion protocol (C02-micro-80, Protifi). Briefly, frozen pellets of 10^7^ cells were thawed on ice in 100 µl 100 mM TEAB buffer (triethylammonium bicarbonate, natural pH 8.5; Supelco 18597) and lysed by probe-sonication on ice-water, using a 1/16” inch microtip and a Fisher Scientific Sonicator, model FB505, with 20% amplitude and 10 cycles of 1 second on, 3 seconds off. An equal volume of 10% SDS in water was mixed and incubated for 55 min at room temperature, followed by a 5 min additional incubation at 55°C. Samples were spun for 30 min at 21,000 x *g* at 4°C and the supernatant (S21) was transferred to a new microtube. 200 µl of each S21 was reduced and alkylated by incubation in 5 mM TCEP (GoldBio) for 15 min at 55°C, then supplemented with methyl methanethiosulfonate (Sigma-Aldrich 208795) to 20 mM followed by a further 10 min incubation at room temperature in the dark. 10 µl, equivalent to ∼5 x 10^5^ cells or ∼50 µg protein, was transferred to a 1.5 mL Protein LoBind tube (Eppendorf) and acidified by vortexing for 30 seconds with 1 µl 27.5% H_3_PO_4_. 165 µl of Binding Buffer (BB: 100 mM TEAB, pH 7.55, 90% MeOH (LC/MS-grade, Fisher A456-4)) was added and the sample was loaded onto an S-Trap micro column (C02-micro-80, Protifi) and spun for 30 s at 4000 x *g*. The column was washed 8 times each with 150 µl BB, the final spin being for 1 min. The columns were transferred to fresh LC/MS-grade MeOH-rinsed tubes. 20 µl of 0.125 mg/ml trypsin (sequencing grade, Promega V511A) in 50 mM TEAB was applied to the column, which was capped and incubated overnight at 37°C in a moist chamber. The S-Trap column was sequentially eluted with 40 µl 50 mM TEAB, 0.2% formic acid (LC/MS-grade, Pierce PI28905) and 0.2% formic acid in 50% acetonitrile (LC/MS-grade, Fisher A955-4). The eluates were pooled, taken to dryness by vacuum centrifugation, and resuspended in 100 µl 0.05% TFA, 5% acetonitrile. 50 µl were dried again, reconstituted in 0.1% heptafluorobutyric acid (Fluka Biochemika 77249), and purified on C18 Zip-tips (Agilent Bond Elut Omix) as described in Boland et al (46). For two 5 h samples, the other 50 µl was dried down, redissolved in 0.1% TFA, and fractionated using the Pierce high pH Reversed-Phase Peptide Fractionation kit (Thermo Scientific, #84868) into 9 pools according to the manufacturer’s instructions. Each were analyzed separately below and together was referred to as a 2D analysis, in contrast to the standard 1D analyses.

### Mass Spectrometry of peptides for proteomic analyzes

Peptides were separated on a C18 nano-column (Thermo Acclaim PepMap 100 C18 series) using an Ultimate 3000 nano-HPLC, and directly infused into a Q-Exactive Plus Orbitrap Mass Spectrometer (Thermo Fisher) as described (46). Raw files were processed in Proteome Discoverer 2.5 using a *D. discoideum* protein database essentially as described in Boland et al. (46) with the following modifications: dynamic oxidation of Met, dynamic loss of N-terminal Met with or without acetylation of N-terminal Met, plus static carbamidomethylation of Cys. Proteome Discoverer 2.5 calculated protein abundances for proteins identified at high (FDR<0.01) and medium confidence (FDR<0.05) were compared with control (t0, 21% O_2_), or same h control samples kept at 21% O_2_ if indicated, using the MetaboAnalyst 5.0 data analysis tool (47). The MS proteomics datasets, which also include proteins identified with low confidence (FDR<0.10), are summarized in Table S6 and are deposited in the ProteomeXchange Consortium via the PRIDE (48) partner repository.

### Proteomic data analyses

Three independent biological replicates, each including 3 technical replicates, were initially performed using the 1D method at each time point. A 0 h, 1 h and 5 h set (Series 1) was performed and analyzed separately from a 0 h, 5 h, 24 h, 1 h reoxygenation and 5 h reoxygenation set (Series 2) owing to differences in the quality and depth of coverage between them. Pearson correlation test was performed on total protein to determine the correlation between time points in both series. The same correlation coefficient was applied to compared total protein and total coding RNA from Table S1B. In addition, a 5 h sample from each series was subjected to 2D analysis, each with 3 technical replicates. These were ratioed to parallel 5 h samples maintained at 21% O_2_ and, owing to differential depth of coverage, were analyzed separately from a combination of the 6 1D 5 h samples (pooled from Series 1 and 2) that was also analyzed relative to 5 h samples maintained at 21% O_2_. 5 h proteins were assigned as differentially expressed if they were quantified based on ≥2 peptides with a p-value <0.05 in at least 2 of the 3 groups if the analysis for the third group was not statistically significant. The 5 h differentially expressed proteins (DEPs) are listed in Table S2. The 24 h DEPs were identified based on ratios relative to t0 that were log2FC≥0.26 or log2FC≤-0.26 (equivalent of FC≥1.2 and FC≤0.83) at p<0.05, and are listed in Table S3. Finally, the total proteins identified from Series 1 and Series 2, along with their relative expression to t0, were merged with the total identified coding RNAs in Table S7. To assist in interpreting the potential functional significance of DEPs at either 5 or 24 h, proteins were manually classified into related functional groups based on experimentation or sequence homologies, with assistance from Dictybase curations. Traditional GO term analysis was not informative owing to the lack of statistical significance given the relatively small number of proteins in each group.

### RT-qPCR protocol

For each experiment, 2 x 10^6^ cells resuspended in 100 µl of HL5 medium were inoculated into 2.9 ml of HL5, previously pre-equilibrated to the desired oxygen concentration using a flow of air/nitrogen mix and maintained at 22 °C. Cells were maintained in a controlled oxygen environment with agitation at 300 rpm. The oxygen concentration was continuously measured using an optical oxygen probe coupled to its oximeter (OXROB3 robust probe and Firesting oximeter, Pyroscience, Aachen, Germany). For the short time course experiment, the HL5 media was pre-equilibrated at 1% O_2_ for 1 h before the cell injection, and collected at 0, 5, 10, 20, 40 and 60 min (n=4). For the drug treatment experiment, cells were exposed to either 1% or 21% O_2_ for 1 h with the different chemicals (n=3). General inhibitors of non-heme dioxygenases were tested at concentrations based on previous studies, including 500 µM cobalt(II) chloride hexahydrate (from a 500 mM stock solution in water, C8661, Sigma-Aldrich, USA), 200 µM deferoxamine mesylate salt (from a 200 mM stock in water, D9533, Sigma-Aldrich), and 1 mM dimethyloxalylglycine (DMOG, from a 100 mM stock in DMSO, 400091, Sigma-Aldrich) (34,49). Complex III of the mitochondrial electron transport chain was inhibited with 37.5 µM antimycin A (from a 5 mM stock in ethanol, A8674, Sigma-Aldrich), and protein synthesis was inhibited using 400 µM cycloheximide (from a 200 mM stock in DMSO, C7698, Sigma-Aldrich) (40,50). Cells were treated just before the experiment, except for cycloheximide which was added 30 min in advance. To test the effect of the oxygen concentration on the hypoxic response, cells were exposed to 1%, 2%, 3%, 5%, 10%, 15% or 21 % O_2_ for 1 h (n=3).

For each experiment, total RNAs were extracted from 2x10^6^ cells (RNeasy Plus Mini Kit, 74134, Qiagen, Germany). Extracted RNAs were quantified and adjusted to 1 µg to synthesize cDNAs (First Strand cDNA Synthesis Kit for RT-PCR, 11483188001, Roche, Switzerland) using random primers (250ng/µl). cDNAs were diluted at 1/20 to perform qPCR (DyNAmo Flash SYBR Green qPCR Kit, F415L, Thermo Fisher Scientific, USA) (Thermocycler BIOER qPCR LINEGENE 9600 PLUS) using the primers following primers: *rnlA*-F (5′-GCACCTCGA TGTCGGCTTAA-3′), *rnlA*-R (5′-CACCCCAACCCTTGGAAACT-3′), *cxgE*-F (5′-ATGTCCCAC GCATTACCAG-3′), *cxgE*-R (5′-CATAGGCAGCGTAAAAATCTTCTC-3′), *cxgS*-F (5′-AAGTTG TTAAATCTCAACTC-3′), *cxgS*-R (5′-TTTATCAACGCCATATTTAA-3′), *fhbB*-F (5′-GATGGT AAAGAGATAGCTAC-3′), and *fhbB*-R (5′-CTGATAAACTGTAATGTCTGAC-3′). For each qPCR, genes were tested in duplicate.

### RT-qPCRs analyses

For each RT-qPCR, the relative gene expression was determined using the 2^-ΔΔCt^ method with *rnlA* as a reference gene (51). For the short time course, the relative expression was calculated using t0 (21% O_2_) as a standard. In the experiment involving chemicals, the relative expression of cells exposed to 1% O_2_ was calculated by using the 21% O_2_ condition with the corresponding chemical as a reference, to assess the effect of hypoxia on gene expression. For cells cultured under 21% O_2_, the condition without the drug was used as the standard to evaluate the effect of the drug under normoxic conditions. For each gene, the percentage of induction at both 21% and 1% O_2_ were calculated using the hypoxic expression level without the drug as the 100% expression. Finally, for the oxygen concentration RT-qPCRs, the relative expression was calculated using the 21% O_2_ condition as the standard. Half-induction (50%) was determined by normalizing the maximum induction, which occurred at either 1% for *rabR*, *srsA* and *fhbB* or 2% O_2_ for *cxgS*, to 100%. The percentage of expression for the other oxygen concentrations was then calculated accordingly. A curve fit using a logarithmic function was used to identify the oxygen concentration corresponding to 50% induction (Fig. S2). For all RT-qPCR experiments, the mean value of the replicates and the standard deviation were calculated.

### RT-qPCR variability analysis

Data from the 10 different experiments corresponding to t0 (21% O_2_ without chemical) and 1 h hypoxia (1% O_2_ without chemical) were combined to assess the main source of variability. The ratios Ct_gene_/Ct_rnlA_ were then calculated. In parallel, TPM at t0 and 1 h hypoxia were also compared, using the triplicates from Table S1A (n=3). For data visualization, TPM values were represented in log2.

## Results

### Hypoxia reduces cell division and triggers two distinct temporal transcriptomic responses

The response of growth stage *Dictyostelium* amoebae residing in nutritive growth medium to acute (1 h), short-term (5 h), and chronic (24 h) hypoxia, followed by reoxygenation (1 h reox and 5 h reox), was examined to cover a time period that at normoxia allows for ∼3 population doublings. Cell density slightly increased between 0 and 5 h, then slowed down further between 5 and 24 h before stabilizing during reoxygenation, suggesting that hypoxia reduces cell cycling and that recovery takes time (Fig. 1A). In contrast, the total RNA content per cell, including polyA RNA and rRNA, strongly increased between 5 and 24 h, indicating a robust transcriptomic response (Fig. 1B). This was confirmed by the turnover rate calculated on a per transcript analysis resulting from the RNA-seq data across time points. We calculated an overall turnover of approximately 20% between each time point, with a marked increase to 40% between 5 and 24 h (Fig. 1C).

**Figure 1:**
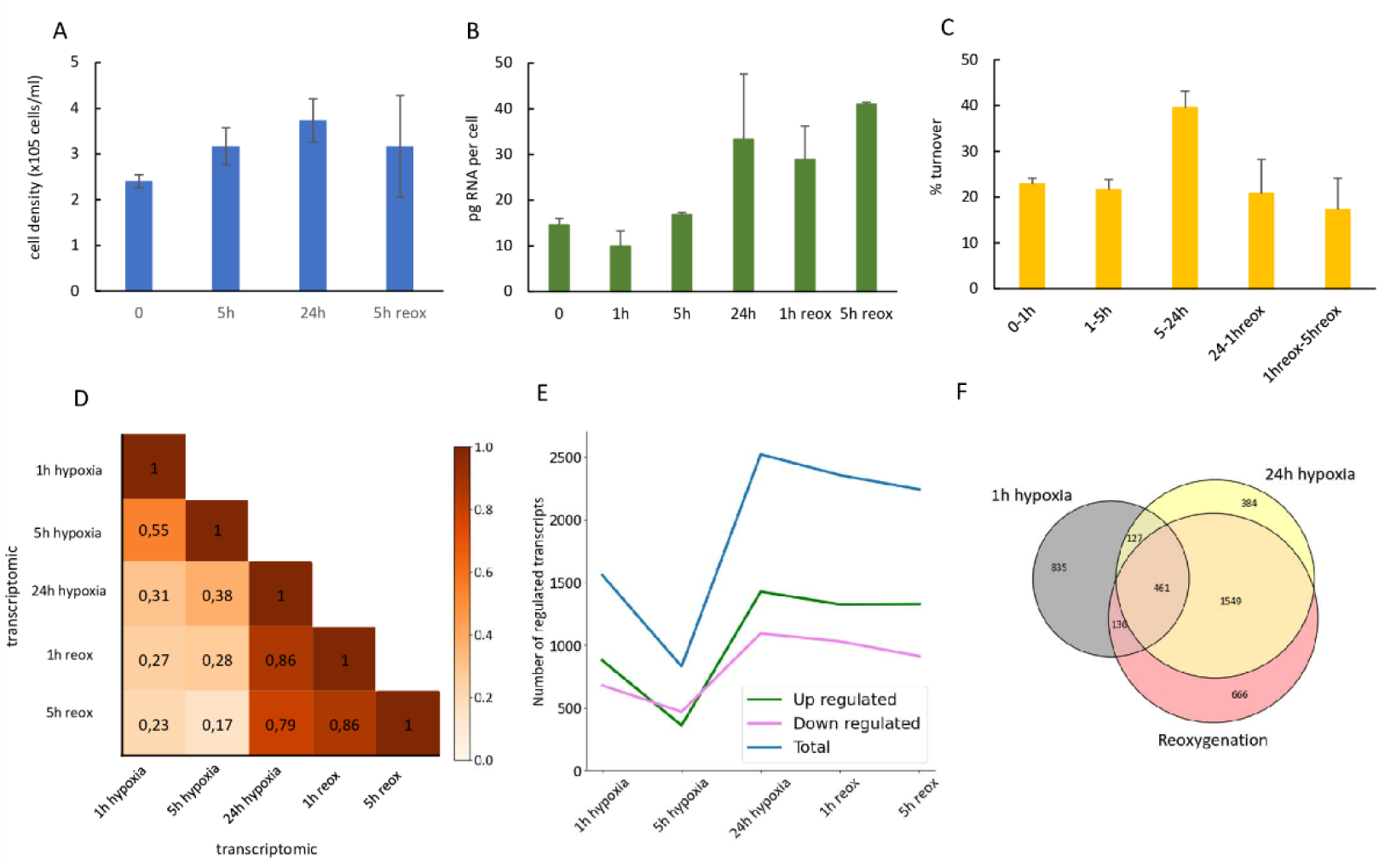
Global transcriptomic response to hypoxia over the time course. **A:** Effect of hypoxia (1% O_2_) on cell division. Cell density was estimated using a hemocytometer. The mean value of the replicates ± the standard deviation is shown on this graph. **B:** Total RNA quantity across the time course. This RNA measurement reflects the total RNA extracted from the collected cells, and was later used for library preparation. The average value of the replicates ± the standard deviation is displayed on this graph. **C:** RNA turnover across the time course. The turnover was calculated by subtracting TPM values between two successive time points for each gene, representing both newly synthesized and degraded RNA. Then the absolute values of all genes TPM differences were summed. Finally, the sum was normalized to a ratio of 1 million and converted into a turnover percentage. The average value of the replicates ± the standard deviation is displayed on this graph. **D:** Correlation between transcripts expression over the time course. Pearson correlation coefficients were calculated between all time points for all coding RNAs (12,265 transcripts). **E:** Number of regulated transcripts at each point of the time course compared to the t0 (21% O_2_). An ad hoc code was applied to remove transcripts with at least one missing expression during the time course. From that list of 11,395 transcripts, only those with a significant differential expression (log2FC≥1 or log2FC≤-1; FDR<0.05) are represented. **F:** Venn diagram representing the overlap among the early, late and reoxygenation responses. Out of the 11,395 transcripts, only those with a significant differential expression (log2FC≥1 or log2FC≤-1; FDR<0.05) at 1 h, 24 h and both reoxygenation time point are shown

Additionally, correlations between each time point of the RNA-seq data were calculated, based on the 12.265 detected transcripts associated with coding genes (summarized in Table S1B). We found that the 1 h response was slightly correlated with the 5 h but not to the 24h response (Fig. 1D). In contrast, the 24 h time point was poorly correlated with earlier time points, but displayed a strong correlation with both reoxygenation time points. The dynamics of only significantly regulated transcripts (log2FC≤1 or log2FC≥1; FDR<0.05) were analyzed to confirm these conclusions, by removing transcripts with at least one missing expression during the time course. Among the resulting 11.395 transcripts, we observed that 32% of the detected coding RNAs were significantly regulated during hypoxia. As shown in Fig. 1E, 1,560 genes were differentially expressed at 1 h compared to t0 (21% O_2_). The number of regulated genes dramatically decreased to around 830 at 5 h, before sharply increasing to 2,531 genes at 24 h following hypoxia. This number remained stable during reoxygenation. These data suggest that the hypoxic response is mediated in two stages, at 1 h and 24 h. This conclusion was further validated by analyzing overlapping responses between those main time points and reoxygenation (Fig. 1F). Specifically, 8% of the genes regulated at 1 h were also regulated at 24 h (127 out of 1,559). In contrast, 61% of the genes regulated at 24 h were also regulated during reoxygenation (1,549 out of 2,521).

Together, these data showed a strong transcriptomic response to hypoxia that occurred in two steps: a rapid adaptation, during the first hour, followed by a later response that persisted during reoxygenation.

### Protein regulation also presents temporal phases with a moderate positive correlation with the transcriptomic data

Parallel samples from the same time course were subjected to a mass spectrometry based proteomics workflow to identify proteins and determine their abundances. (see Methods), We then evaluated the conservation of the 1,762 proteins that were detected at sufficient levels to be quantified of total protein regulation between time points. Changes recorded at 1 h were infrequently conserved at 5 h, with a weak positive correlation of 0.37 (Fig. 2A). This might reflect an early phase of adaptation but, given the limited time for protein levels to change and the smaller number of affected proteins compared to 5 h, the 1 h data are not systematically summarized. A moderate positive correlation of 0.43 was calculated between 5 h and 24 h time points (Fig. 2B). The data suggest discrete early and late stages of adaptation that are, however, not as independent as suggested by the transcriptomic data. In contrast, a strong positive correlation between 24 h and reoxygenation time points ranging from 0.81 to 0.91 was observed. This indicated delayed proteomic recovery during the 5 h of reoxygenation, as also observed transcriptomically. The strong similarities between the 24 h and 1 h of reoxygenation data validate the assignments of differentially expressed late phase proteins described below.

**Figure 2:**
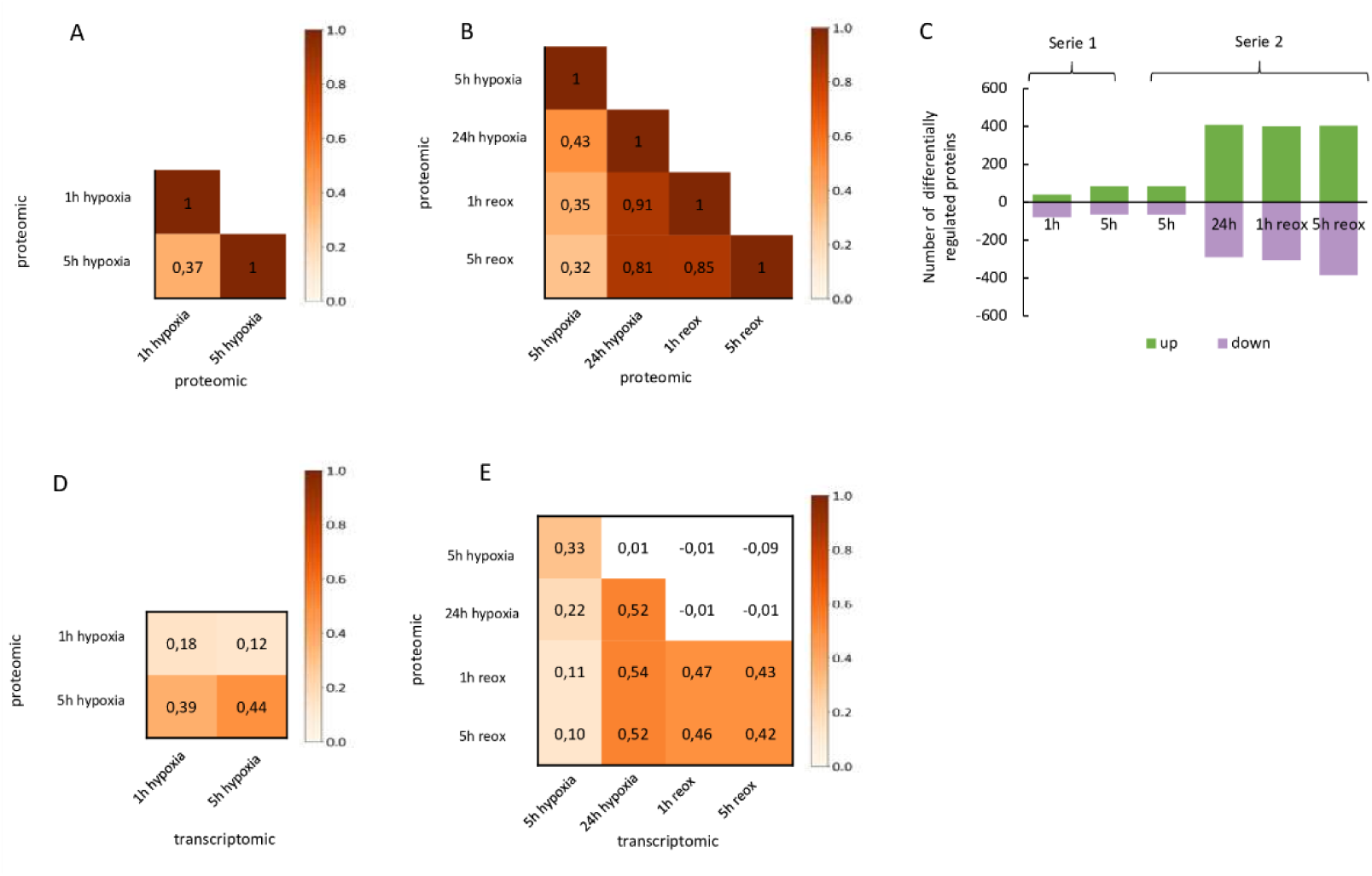
Global proteomic response to hypoxia over the time course and correlation with the transcriptomic data. Only proteins detected with medium or high detection confidence and quantitated based on ≥2 peptides were analyzed. **A:** Correlation at the proteomic level across Series 1 (early response). Pearson correlation coefficients were calculated between the 1 h and 5 h hypoxia time points. **B:** Correlation at the proteomic level across Series 2 (late response). Pearson correlation coefficients were calculated between the 5 h, 24 h and both reoxygenation time points. **C:** Number of regulated proteins at each time point of the time course compared to the t0. Only proteins with significant differential expression (log2FC≥0.26 or log2FC≤-0.26; p<0.05) are shown. **D:** Correlation between transcriptomic and proteomic data across Series 1 (early response). Pearson correlation coefficients were calculated between the 1 h and 5 h hypoxia time points. **E:** Correlation between transcriptomic and proteomic data across Series 2 (late response). Coefficients were calculated between the 5 h, 24 h and both reoxygenation time points.

Additionally, the dynamics of significantly regulated proteins (log2FC≥0.26 or log2FC≤- 0.26 ; FDR<0.05) were analyzed. Out the 1,762 detected proteins, 2.3%, 4.8% and 23% of the total were differentially increased at 1, 5, and 24 h, respectively, whereas, 4.8%, 3.7% and 17% were decreased (Fig. 2C). Thus by 24 h, 40% of the proteome quantitated was affected by treatment at 1% O_2_, indicative of major proteomic remodeling during hypoxia. Moreover, the approximate quintupling of differentially expressed proteins at 24 h of adaptation corresponds to the burst differentially expressed transcripts.

To assess potential linked regulation, the correlation between the transcriptomic and proteomic datasets was calculated. While only a weak correlation coefficient of ∼0.18 was observed at 1 h (Fig. 2D), it improved to 0.33 at 5 h and ∼0.5 at 24 h and during reoxygenation (Fig. 2E). The correlations at the later time points indicated that a substantial fraction of the proteomic changes could be explained by changes in transcript levels, and lower correlations at earlier time points are consistent with greater time required for proteomic responses. This is supported by the fact that the 1 h transcriptomic data show a stronger correlation with the 5 h proteomic data than with the 1 h proteomic data.

Taken together, the data indicate that the proteomic response is also temporally regulated, with a discrete early regulation phase that shows limited correlation with the transcriptomic response. In contrast, the later proteomic regulation, which persists during reoxygenation, is more closely aligned with the transcriptomic response. Therefore, we conducted further analyses to better understand the biological processes associated with these two distinct responses at both transcriptomic and proteomic level.

### The early transcriptome response (1 h) is associated with ribosome biogenesis

We first investigated biological functions associated with the early response to hypoxia. 677 transcripts were exclusively regulated after 1 h of hypoxia, with a majority being weakly up-regulated compared to t0 (Fig. S3A). No significant biological function was identified for the down-regulated transcripts. The proteins encoded by those up-regulated transcripts were identified as forming a single cluster of strongly interacting proteins (Fig. 3A), suggesting participation in the same pathway. We showed that these transcripts were enriched relative to their representation in the total genome across multiple steps of the ribosome biogenesis pathway (Fig. 3B). These effects, which included both nuclear and cytoplasmic processes, were validated by fold enrichments that showed an increase in genes linked to both rRNA and tRNA methylation (Table S5A). Uridine modifications were also up-regulated, with notable enrichment in transcripts associated with pseudouridine synthesis and wobble uridine modification in tRNAs. Furthermore, a particularly strong enrichment of genes involved in ribosomal subunit maturation and ribosome assembly was observed. Genes encoding enzymes for the histidine-to-diphthamide modification, a key modification required for translation, were also over-represented. Finally, genes involved in rRNA transcription were up- regulated, alongside positive regulation of RNA polymerase I.

**Figure 3:**
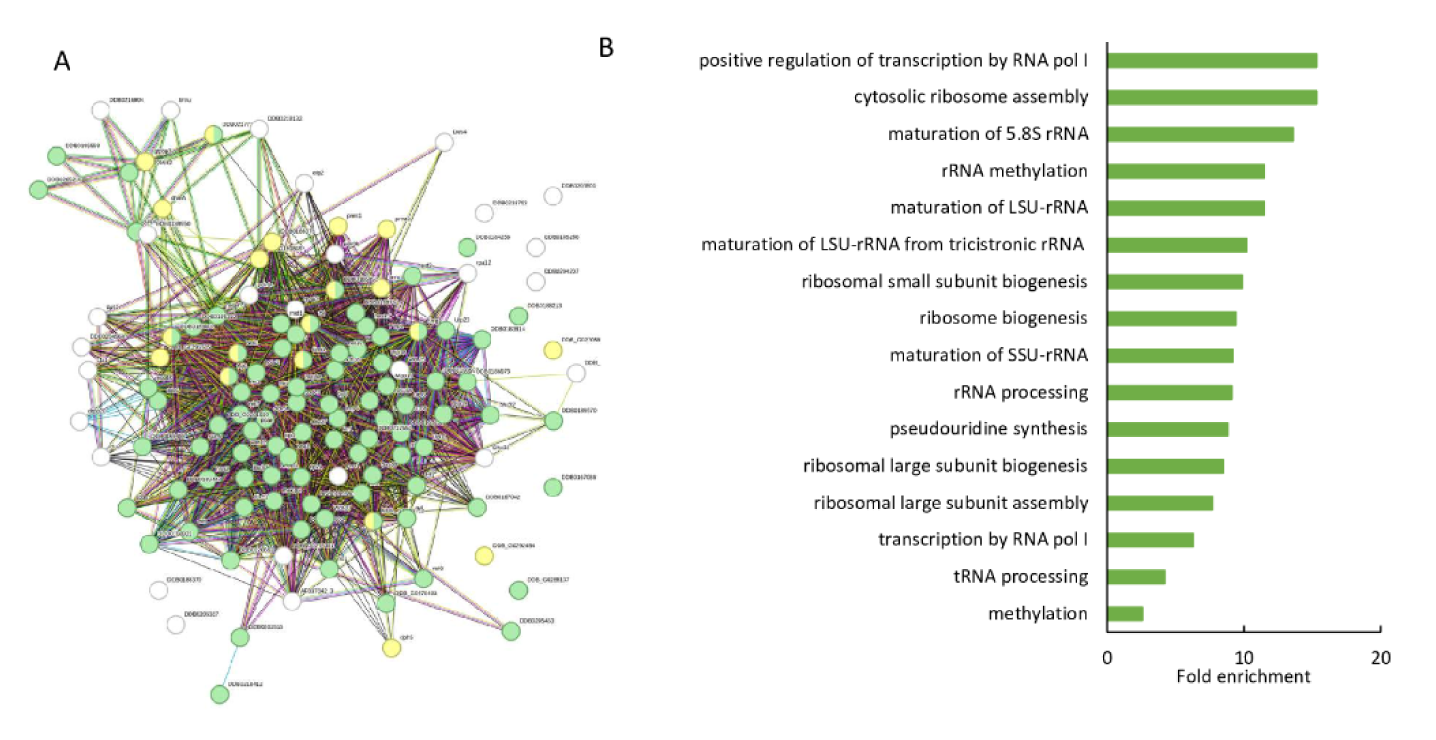
Acute hypoxia induces the up-regulation of transcripts related to ribosome biogenesis. **A:** Interactions between proteins encoded by transcripts up-regulated (log2FC≥1; FDR<0.05) after 1 h of hypoxia were evaluated using STRING analysis. Proteins involved in ribosome biogenesis are represented in green and those involved in total methylation, including rRNA methylation, are in yellow. **B:** Fold enrichment of transcripts up-regulated at 1h after hypoxia. Only up-regulated transcripts at 1 h were analyzed (log2FC≥1; FDR<0.05), and only significant fold enrichments (FDR<0.05) are shown (from Table S8A). The fold enrichment was provided by DAVID comparing the representation of the GO category transcripts in the set of differentially expressed transcripts relative to the 7539 genes from *Dictyostelium* reference genome in the DAVID database.

Collectively, the transcriptomic data indicate the potential for rapidly induced ribosome biogenesis following hypoxia. Moreover, mid-hypoxia does not have a distinct biological function of its own but rather extends the acute hypoxic response (Table S8B). These findings suggest that mid-hypoxia represents a transitional phase between rapid adaptation and the slower, long-term response.

### The early proteomic response (5 h) is mostly associated with an increase of O2-related proteins and lipid metabolism

We reasoned that the early transcriptome response would be most likely reflected in the 5 h proteome. To improve sensitivity, two of the six 5 h samples were also analyzed by the 2D method. The combined analysis of all 5 h 1D and 2D data increased the number of quantified proteins by about 50% to 2,743 proteins. Of the 131 proteins that were induced in this expanded list (Table S2A), 40% were associated with O_2_ as O_2_-binding proteins, re-dox enzymes, or enzymes utilizing O_2_ as a substrate (Fig. 4A). In contrast, only 11% of the down- regulated proteins were associated with this group. Similarly, proteins associated with lipid metabolism, central carbon and amino acid metabolism, signaling, cytoskeleton, and Ca^2+^, were much more highly enriched in the up-regulated group compared to the down-regulated group. A notable example of a coordinated transcriptional and proteomic response was the dihydrosphingosine desaturase DesA associated with lipid metabolism. This gene is induced at the transcriptomic level as early as 1 hour after hypoxia and shows an increase in protein abundance after 5 h (Table S7). For comparison and reference, Table S4A lists the remaining proteins for which no significant abundance changes were observed; Tables S4C and S4B list proteins identified by only one peptide and at low confidence with ≥2 peptides respectively, which were not analyzed.

**Figure 4:**
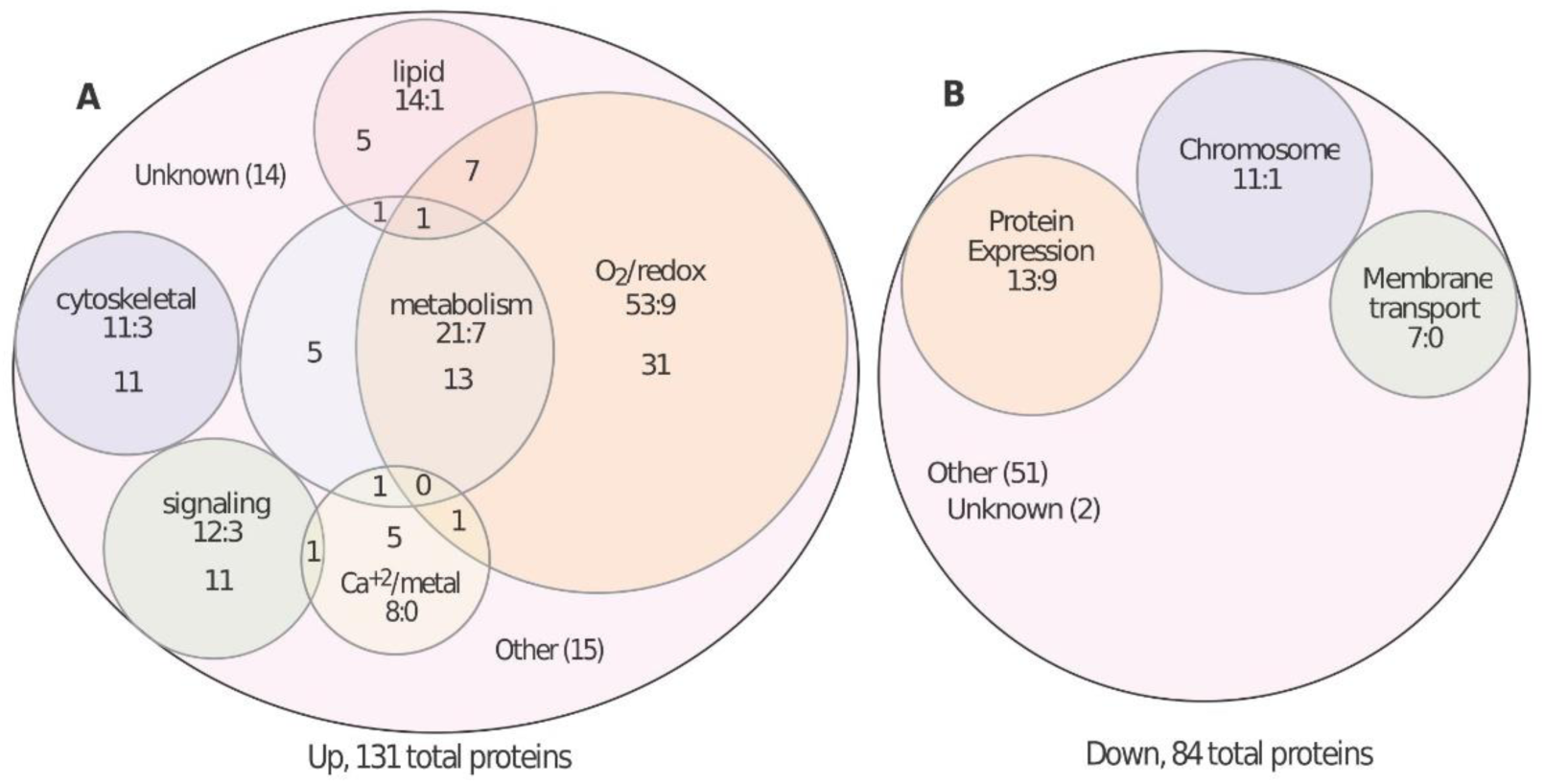
Classification of differentially expressed proteins (DEP) after 5 h at low O_2_. Within the outer circles, proteins related by sequence or functional annotations, and that were highly enriched in either up-regulated on down-regulated lists (Table S2), are identified by circles scaled to group size. Under the group labels are the number of proteins in the up or down list, followed by the number of proteins in the opposite (down or up) list. Proteins assigned to multiple groups are represented in the overlap regions. **A:** Classification of up-regulated proteins after 5 h at 1% O_2._ Proteins that were increased (log2FC≥0.26; p<0.05) from the 3097 proteins that were quantified with ≥2 peptides in the amalgamated 1D and 2D data, are represented by the outer circle. **B:** Classification of down-regulated proteins after 5 h at 1% O_2._ Same for proteins that were down-regulated (log2FC≤-0.26; p<0.05).

In comparison, the 84 proteins that were down-regulated (Table S2B) were more highly enriched with functions such as with chromosomes, protein expression, and membrane transport (Fig. 4B). Notably, the steady state levels of the great majority of ribosome-related proteins were not affected, and those that were affected were more likely to be decreased than increased. This inconsistency suggests that the effects of up-regulation of ribosome-related transcripts at 1 h were transient or supporting increased protein turnover rather than abundance.

### The long-term response is related to multiple processes that persist during reoxygenation

We then analyzed the response to chronic hypoxia, which largely persists during reoxygenation, as identified in both transcriptomic and proteomic data. In the transcriptome, 986 transcripts were regulated at all these time points, and a significant portion of them were overexpressed (Fig. S3B). In contrast to the short-term response, the proteins associated with these up-regulated transcripts are less interconnected, suggesting that distinct functions are involved (Fig. 5A). In comparison, about 700 of the ∼1,700 proteins that were quantified were differentially expressed at 24 h. All transcripts and proteins that are not documented in the figures can be found in Table S7. Additionally, the differentially expressed proteins are listed in Table S3, where they are assigned to groups based on function or sequences suggestive of function. The expression trajectories from 5 h of hypoxia through 5 h of reoxygenation are plotted for some examples (see below). For comparison and accessibility, Table S5 lists 24 h proteins for which there was no evidence for differential expression (Table S5A), as well as proteins detected on the basis of a single peptide (Table S5C) or at low confidence though with ≥2 peptides (Table S5B).

**Figure 5:**
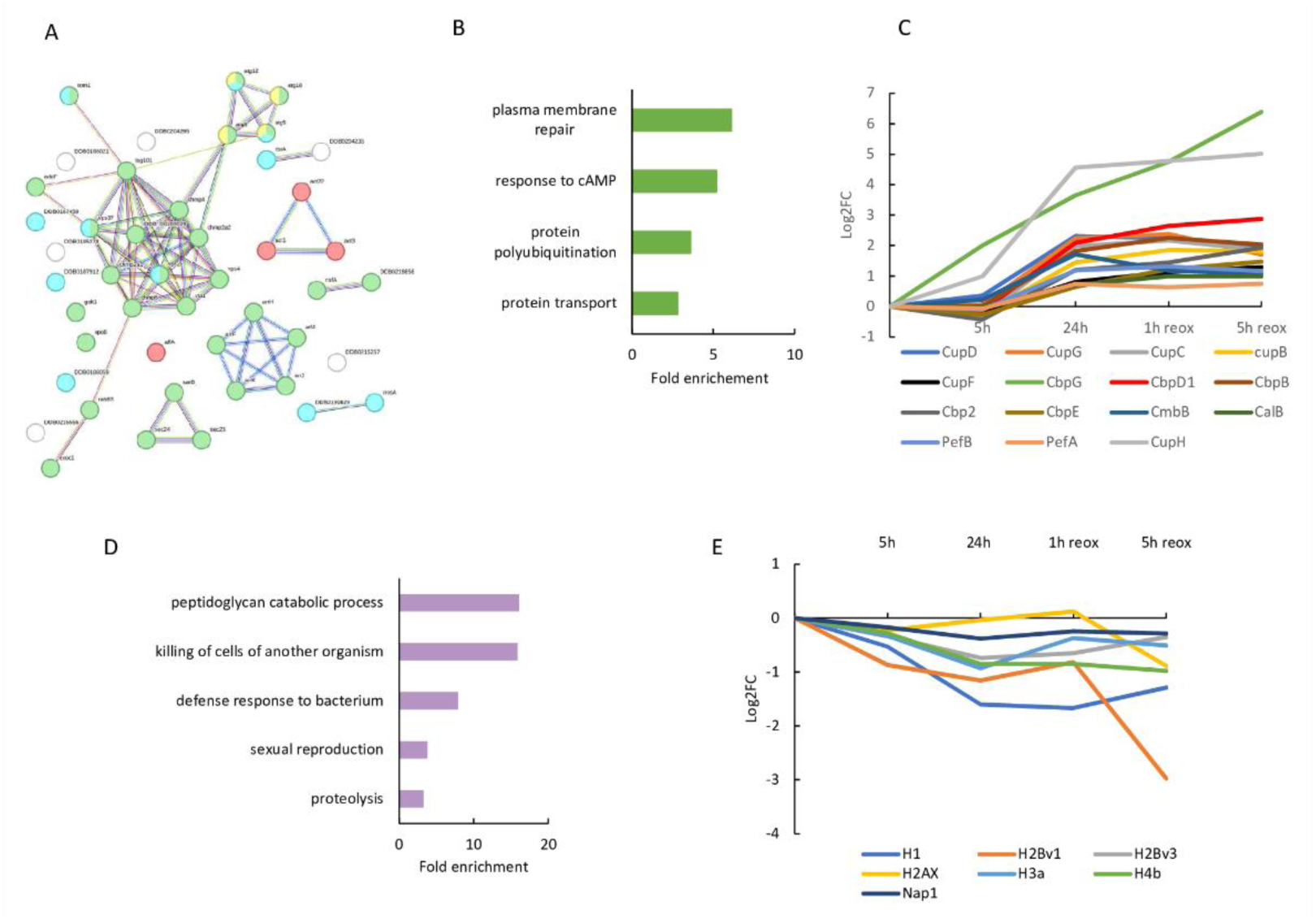
Chronic hypoxia regulated multiples processes that persists during reoxygenation. **A:** Interactions between proteins encoded by transcripts up-regulated at 24 h and during reoxygenation. Interactions were evaluated using STRING analysis for up-regulated transcripts (log2FC≥1, FDR<0.05). Proteins involved in protein transport are represented in green, those in the ubiquitin catabolic process in blue, those responding to cAMP in red, and those associated with autophagy organization in yellow. **B**: Fold enrichment of transcripts up-regulated after 24 h hypoxia and during reoxygenation. Only up-regulated transcripts were analyzed (log2FC≥1; FDR<0.05), and only significant fold enrichments (FDR<0.05) are shown (from Table S8C). **C:** Calcium associated proteins regulated at 24 h and during reoxygenation. All significantly regulated proteins (p-value>0.05) are shown. **D:** Fold enrichment of down-regulated transcripts after 24 h hypoxia and during reoxygenation. Only down-regulated transcripts were analyzed (log2FC≤-1; FDR<0.05), and only significant fold enrichments (FDR<0.05) are shown (from Table S8C). **E:** Regulation of histones and histone-related proteins during chronic hypoxia and reoxygenation. Note that H2Bv1 and H3a were low-confidence hits.

An enrichment was observed for transcripts involved in protein transport, including several Vacuolar Protein Sorting genes (Fig. 5B, Table S8C). Among them Vta1, Vps4, Vps60, Vps32 are similarly up-regulated at the protein level at 24 h. Another transcript related to the ESCRT-I complex, *tsg101*, was also up-regulated, along with its associated transcript *tom1*. Within the protein transport category, many transcripts associated with small GTPases were strongly induced, such as *arrF*, *arrK*, *arrH*, and *arrJ*, with *arfA* showing more moderate up- regulation. As noted below, many transcripts and proteins associated with the cytoskeleton, and which might synergize with protein transport, were also up-regulated. Regarding potential regulation of these processes, many transcripts and 15 proteins associated with calcium (Table S1D, Fig. 5C) were strongly over-expressed during chronic hypoxia. This includes calcium up- regulated proteins (CUPs), Ca^2+^-binding proteins, and calmodulin-related proteins. In the transcriptome, the CUPs exhibited a distinct pattern, with induction at 1 h and decreased expression at 5 h, followed by intense overexpression during chronic hypoxia and reoxygenation. This pattern was especially observed for *cupB*, *cupG*, *cupC* and *cupF*. Though the significance is not clear, dramatic transcript increases for the highly abundant discoidin proteins encoded by *dscA* and *dscD* were observed after 24 hours hypoxia, while dscE increased only after 5 h reoxygenation, and *dscC* expression mostly remained constant over the time course. These findings were confirmed at the protein level for DscA, DscC, and DscE.

Regarding protein degradation, transcripts related to autophagy, including *atg8*, *atg12*, and *atg18*, were upregulated. The expression of *atg5* was also moderately induced, and the proteomic analysis showed that Atg5 as well as Atg8a were strongly represented after 24 h of hypoxia and during reoxygenation (Table S8C). Another enrichment was observed for transcripts involved in protein polyubiquitination, including *nosA* and *rbrA* E3 ubiquitin protein ligases. Even while they were not annotated in the GO biological process, an induction of several ubiquitin-related transcripts, including *ubqJ* and *ubqH* (as well as *ubqK*, *ubqI*, *ubqG*, and *ubqA*) was also observed (Table S1B). In the proteome, 15 of the 17 proteasome-associated, and all 12 of the ubiquitin-associated proteins were differentially up-regulated (Fig. S4A Table S3I). Among them, the proteins UbcB and CulE, were increased proteomically though did not meet the transcriptome criteria for change.

Finally, many glycolytic enzymes were up-regulated at the proteomic level while remaining minimally changed in the transcriptome. The stronger up-regulation of FBPase and PEP-carboxykinase relative to the weaker up-regulation of PFK1 suggested that gluconeogenesis was activated (Fig. S4B, Table S3G). This is consistent with the strong up- regulation of the peroxisome-associated citrate synthase GltA likely involved in the glyoxylate cycle (52), and the up-regulation of acyl-CoA dehydrogenase, both of which might provide carbon for gluconeogenesis (Fig. S4C). However, another enzyme of the glyoxylate cycle, malate synthase, was down-regulated. The TCA cycle did not appear to be affected, as increases in mitochondrial citrate synthase and fumarase was matched by a decrease in isocitrate dehydrogenase (NAD+ form). Glycogen is another potential source of carbon based on the strong up-regulation of Tf2, a putative transcription factor for GlpD (53) . Though Tf2 did not pass our threshold, the *glpD* transcript was strongly increased, whereas the transcript level of *glpV*, the vegetative isoform, was unaffected. Finally, a substantial number of proteins related to oxygen or redox reactions continued to be elevated at 24 h, and outnumbered those downregulated by a 3:1 ratio (Table S3E). However, they comprised only ∼15% rather than 40% of total upregulated proteins at the respective time points.

Among the down-regulated transcripts, a significant fold enrichment was observed for genes involved in the peptidoglycan catabolic process, including lysosome-related genes such as *alyC*, *alyB*, and *lyEh3* (Fig. 5D, Table S8C). This enrichment is associated with the defense response to bacterium and killing of cells against of other organisms. In the proteomic data, 10 proteins associated with the membrane of lysosomes, including the vacuolar ATPase, were reduced in level, and LyEh3 was also reduced though failed meet our cut-off criteria (Table S3F). Transcripts involved in vesicular proteolysis were also down-regulated, with moderate but statistically significant fold enrichment. Within this cluster, serine and cysteine proteases as well as peptidases like *tpp1C* and *tpp1B* were identified. The cluster also included lysosomal cathepsins, specifically *ctsZ* and *ctsD and cprD*, that were also strongly down regulated at the protein level (Table S3F). Moreover, proteomic analyses highlighted that proteins associated with protein synthesis were largely depressed during chronic hypoxia and reoxygenation (Table S3L), though these changes were not anticipated from the transcriptomic analyses. Finally, an enrichment was observed for sexual reproduction for down-regulated transcripts (Fig. 5D). However, this annotation was based on the biological aspect of an ancestor (IBA) and lack any additional characterization (Table S8C).

At the protein level, all 53 of the cytoplasmic ribosomal proteins that were quantified were reduced to a statistically significant extent (Table S3K). Similarly, 4 of the 5 detected mitochondrial ribosome subunits were similarly reduced, though one (Mrpl13) was increased. These effects were mirrored by the reduction in level of all 7 cytoplasmic tRNA synthases/ligases that were detected (Fig. S4D, Table S3J); the one mitochondrial tRNA synthase detected (54) was increased. These effects, except for the mitochondrial tRNA synthase, persisted through the 5 h reoxygenation period, implying continued reduced translational activity. Capacity for protein secretion appeared diminished based on decreased levels of all 6 rER enzymes quantified, and increased levels of all 4 COPII proteins associated with retrotranslocation back to the rER (Table S3F). The majority (14/17) of other factors associated with protein translation were also decreased, with the exceptions potentially supporting selective translation of a subset of mRNAs. As indicated above, reduced capacity for protein synthesis was matched by increased protein degradation capacity. Finally, proteomic data also showed that many histones (H1, H2Bv1, H2Bv3, H3a, H4b) were moderately repressed at 24 h, but the greatest effect was noted for H1 (log2FC=-1.6) (Fig. 5E, Table S3H). An exception was H2AX, one of two H2a isomers, which was not affected. Interestingly, the transcripts for these histones, except for *H4b*, remained stable.

Taken together, the data imply that chronic hypoxia influences multiple processes, including decreased capacity for protein synthesis and up-regulation of proteasomal degradation and protein transport, and catabolic lysosomal activities that might also connect with increased autophagy. Proteomic analyses revealed that chronic hypoxia was also associated with an increase in the activity of central carbon metabolism though the precise direction of metabolic flow was unclear. Interestingly, most changes in levels of metabolic enzymes and proteins involved in anabolic processes were not detected at the transcriptomic level. Thus, increased proteolysis might explain decreased protein synthesis capacity, but other mechanisms such as translational regulation may need to be invoked to explain effects on metabolic enzymes.

### Chronic hypoxia specifically down-regulates genes related to cell division

Among the transcripts regulated in the long-term response, some returned to their basal state immediately upon reoxygenation. Most were repressed (Fig. S3C), with no functional clusters identified for the up-regulated transcripts. The down regulated transcripts are associated with cell division, with significant fold enrichments for DNA replication and mitotic division (Fig. 6A and B, TableS8D). More specifically, the *orcD* transcripts, which encode a subunit of the origin recognition complex and *cdc45*, were downregulated. At the proteomic level, PCNA, Rfa1, PolD3, and PolD2 (the latter two identified based on a single peptide) were found to be repressed (Fig. 6C). In contrast, RcbA supposedly involved in DNA repair was showed to be induced. The expression of transcripts related to the mitotic spindle assembly checkpoint, including *anapc7* and *bub1*, were down-regulated. A few repressed transcripts involved in chromosome separation were manually identified as proper annotation were lacking. These include the inner centrosome *icpA*, the kinetochore *spc25* and the kinesins *kif12* and *kif13*. Consistently, two main proteins that contribute to cell cycle progression, Cdk1 and Gnl3, were down regulated at the protein level but did not meet the cut-off for transcript change. This is consistent with regulation via protein turnover rather than synthesis, as occurs for other regulators of cell cycle progression.

**Figure 6:**
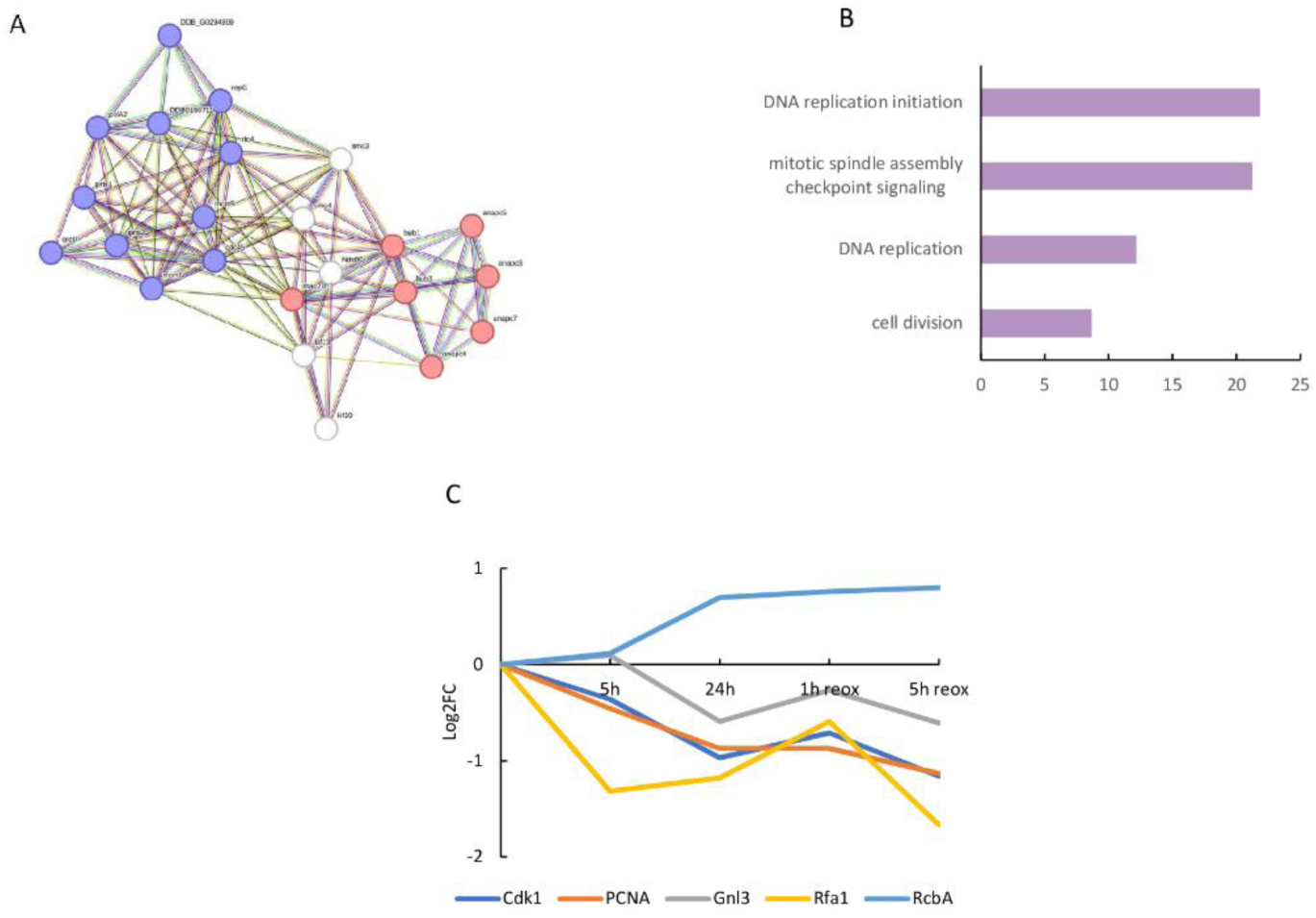
Chronic hypoxia represses the expression of genes related to cell division. **A:** Interactions between proteins encoded by transcripts down-regulated (log2FC≤-1; FDR<0.05) after 24 h of hypoxia. These direct and indirect interactions were evaluated using STRING. In purple are represented the proteins involved in DNA replication are represented in purple; proteins involved in the mitotic spindle are in red. **B:** Fold enrichment of transcripts down regulated (log2FC≤-1; FDR<0.05) after 24 h of hypoxia. Only significant fold enrichments (FDR<0.05) are shown (from Table S8D). **C:** Regulation of cell cycle-associated proteins. Only significantly regulated proteins (log2FC≥0.26 or log2FC≤-0.26; p-value>0.05) are shown.

Collectively, these results indicate that hypoxia inhibits cell division, likely by limiting both DNA replication and the mitotic metaphase-to-anaphase transition. This response takes time to set up but disappears quickly when oxygen concentrations return to normoxia. These findings align with the near arrest of cell division between 5 and 24 h (Fig. 1A), despite an expected 8-fold increase in cell number after 24 h in normoxia.

### Chronic hypoxia stably induces genes linked to cAMP-dependent aggregation

We surprisingly found that transcripts related to the cAMP pathway were induced after 24 h of hypoxia and during reoxygenation (Fig. 7) This well-studied pathway contributes at many levels to starvation-induced development, so further characterization was performed (Table S1E). Manual analysis was required as many components associated with chemotaxis are known only in *Dictyostelium* and poorly included in global databases. Its 3 adenylate cyclases were up-regulated in the 24 h transcriptome, with *acaA* continuing to increase during reoxygenation (Fig. 7A). The cAMP receptors *carC* and *carD* were repressed during acute hypoxia, and then moderately up-regulated during chronic hypoxia and reoxygenation (Fig. 7B). *CarA* was strongly repressed at 1 h after hypoxia, before turning to be slightly induced at 24 hours and beyond. The induction of the extracellular phosphodiesterase *pdeE* started during chronic hypoxia, while *pde4* and *pde7* expression were mainly correlated to reoxygenation (Fig. 7E). The phosphodiesterase inhibitor *pdiA* was strongly repressed during acute hypoxia, before being up-regulated during chronic hypoxia and reoxygenation. The intracellular cAMP phosphodiesterase RegA was induced at 5 h reoxygenation, while the associated histidine kinase genes *dhkC* and *dhkB* were induced by 24 h (Fig. 7C). Finally, members of the cAMP dependent kinase pathway, including *pkaR* and *pkaC* as well as a downstream target, the transcription factor *bzpF*, were also induced ((Fig. 7D). Proteins related to those transcripts were not detected, except RegA that showed a strong induction at 24 h but detected with an FDR<0.1 by a single peptide. Overall, the transcriptomic profiles suggested initial hypoxic refractoriness to cAMP until persistent activation of the network at the chronic stage and during reoxygenation.

**Figure 7:**
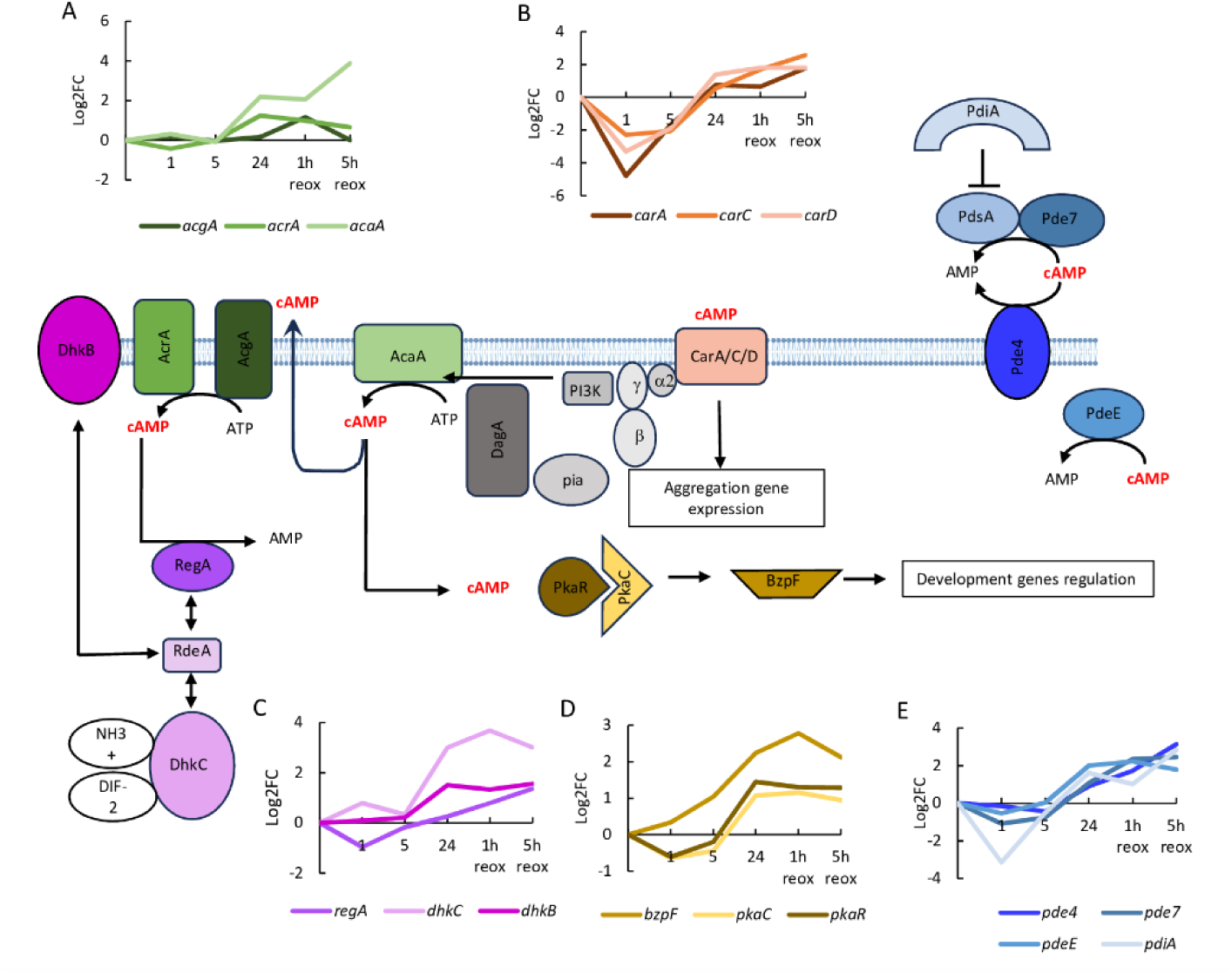
The cAMP aggregation pathway is activated at the transcriptomic level during chronic hypoxia and reoxygenation. Only differentially up-regulated (log2FC≥1; FDR<0.05) transcripts after 24 h of hypoxia or for at least one time point during reoxygenation were plotted. **A**: Transcriptomic profiles of the cAMP receptors. CarA, CarC, and CarD, were differentially expressed. **B**: Regulation of the adenylate cyclases involved in cAMP production. **C**: Regulation of the RdeA/RegA two-component system and their regulatory kinases, DhkB and DhkC. RdeA expression is not represented as it was not differentially expressed. **D**: Transcriptomic profile of the regulatory (PkaR) and catalytic (PkaC) subunits of the cAMP dependent protein kinase A and its associated transcription factor BzpF. **E**: Transcriptomic profile of cAMP phosphodiesterases and an associated inhibitor involved in cAMP degradation.

Moreover, many transcripts and proteins associated with the cytoskeleton were overexpressed during chronic hypoxia and reoxygenation (Table S1D, Table S3D, Table S8C). Many actin-coding transcripts were strongly induced and, at the protein level, the Act21 family (17 different genes with identical protein sequences) and Act3 were also up-regulated. Transcripts encoding actin-binding proteins were significantly up-regulated, with a strong induction of ponticulin transcripts including *ponC* and *ponF*, alongside a more moderate induction of fimbrin transcripts like *fimA*, which also showed increased expression at the protein level. Additionally, some regulators of the actin cytoskeleton, including the *arfGAPs*, were induced at the transcriptomic level. The myosin transcripts *myoC*, *myoK* and *myoB* were induced after 24 h in hypoxia and during the first hour of reoxygenation. At the protein level, MyoK (but not MyoB) and 4 other myosin heavy or light chains were induced. In contrast, a few transcripts related to actin and its organization were down-regulated, including *ponJ*, *ponD* or *act32*. Oxygen stress did not appear to substantially regulate microtubules, though the putative microtubule affinity-Regulating Kinase *mrkB* transcript was strongly induced and proteomic data revealed an up-regulation of the tubulin TubA.

Finally, transcripts and proteins related to cell adhesion were also overexpressed after 24 hours of hypoxia and during reoxygenation (Table S1F, Table S3F). The contact site B transcripts, such as *csbC*, *csbB* and *csbA,* were strongly expressed. The calcium-dependent adhesion transcript *cad2* was also greatly induced, while *cadA* expression remained unchanged. *cad3* was only up-regulated at 5 h of reoxygenation, but with a high level of induction. At the protein level, CsbC was also increased, as were CsbB, Cad2 and Cad3 though they did not meet our cut-off filters. The tiger transcript *tgrB2* was also greatly induced by long hypoxic exposure. Moreover, *sibD* transcripts showed a strong induction under chronic hypoxic conditions, while *sibB* expression increased progressively. Thus, there was evidence for dynamic regulation of intercellular contacts which, together with increased cAMP signaling.

Taken together, these data suggest that chronic hypoxia induces the transcription of genes involved in the cAMP pathway used for chemotactic aggregation including those responsible for cAMP synthesis, reception, degradation, and signal transduction. Additionally, transcripts involved in later stages of aggregation, including those related to actin remodeling and cell adhesion, were up-regulated and identified in both transcriptomic and proteomic analyses. These inductions occur despite the fact that the cells are in an environment not conductive for aggregation as they are maintained in suspension at low density by shaking in growing media.

### Identification of marker genes and potential regulators of the hypoxic response

We surveyed individual transcripts to nominate potential regulators of the hypoxic response and predict marker genes for analysis of the dynamics and mechanisms of the response. The most substantially induced transcript under hypoxia was the small GTPase *rabR*, which exhibited more than a 250-fold increase in expression. This induction was specific to hypoxia, with a return to the basal expression level after 1 h in reoxygenation (Fig. 8A). The starvation-responsive small protein A, encoded by *srsA*, was also strongly and quickly induced by hypoxia, before being repressed during reoxygenation. Both SrsA and RabR were also strongly induced at the protein level, though not surprisingly their changes tended to lag behind that of their transcripts.

**Figure 8:**
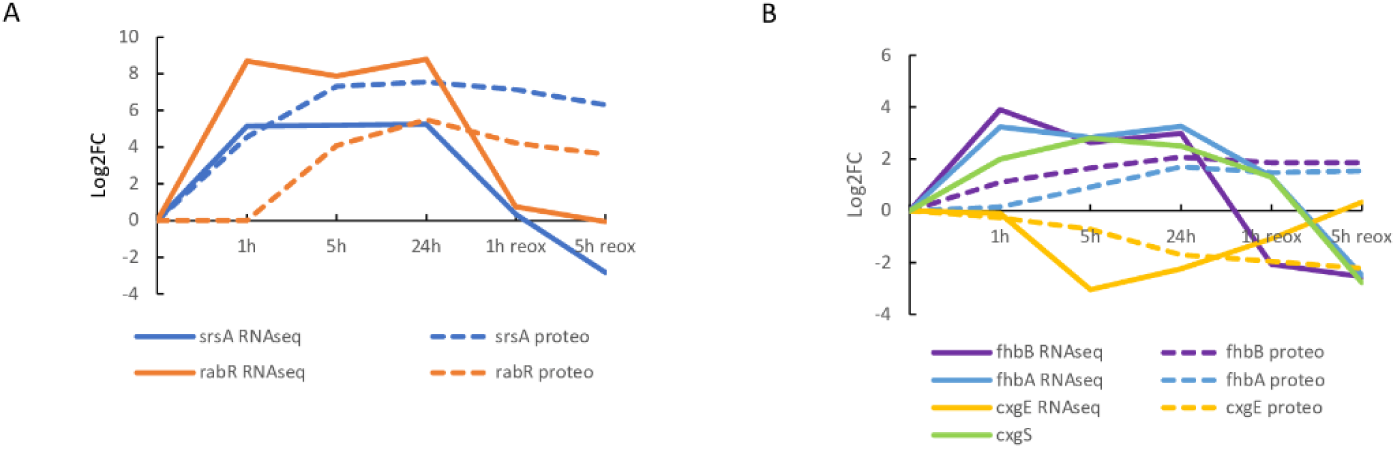
Identification of marker genes and potential regulators of the hypoxic response. Transcriptomic and proteomic regulation of genes in response to hypoxia and reoxygenation. Transcriptomic (from RNA-seq analyses) and proteomic profiles are represented by continuous and dashed lines, respectively. For the 5 h time point, the averages of series 1 and series 2 data are represented. **A:** Regulation pattern of two strongly hypoxia-induced genes used as markers. **B:** Regulation of genes previously described as hypoxia-regulated. CxgS was not detected in the proteomic analyses, likely due to the small size of the protein.

A similar pattern was found for genes already described as induced by hypoxia (39,40). Indeed, the 2 flavohemoglobins, *fhbA* and *fhbB*, were induced by 1 h in hypoxia, but with a lower fold change than *rabR* and *srsA* (Fig. 8B). Additionally, the two genes encoding subunit VII of the cytochrome C oxidase, *cxgS* and *cxgE*, were analyzed. *cxgS* was induced during hypoxia, before being repressed at 5 h of reoxygenation. In contrast, *cxgE* was repressed at 5 and 24 h, before returning to its basal level after the return to normoxia. FhbB, FhbA and CxgE were similarly affected at the protein level with a delay as expected (Fig. 8A and B).

These findings highlight key marker genes and potential regulators that were further studied by RT-qPCR to better understand the hypoxic response and to elucidate regulatory mechanisms.

### Marker genes displays multiple dynamics mechanisms of expression

We further examined the dynamics and molecular mechanisms of transcript expression of the marker genes *srsA*, *rabR*, *fhbB*, and *cxgS*, which represent a diversity of biological functions. Using RT-qPCR, *srsA* and *rabR* exhibited stronger induction after 1 h at 1% O_2_ than *cxgS* and *fhbB*, confirming the RNA-seq findings (Fig. 9A and B). Both *fhbB* and *cxgS* exhibited linear induction patterns, with *fhbB* expression plateauing at 40 min and *cxgS* induction continuing to increase past 60 min as indicated by RNA-seq data (Table S1A). On the contrary, both *rabR* and *srsA* showed strong and almost complete induction within 10 min, while SrsA expression continued to slightly rise over the next 60 min.

**Figure 9:**
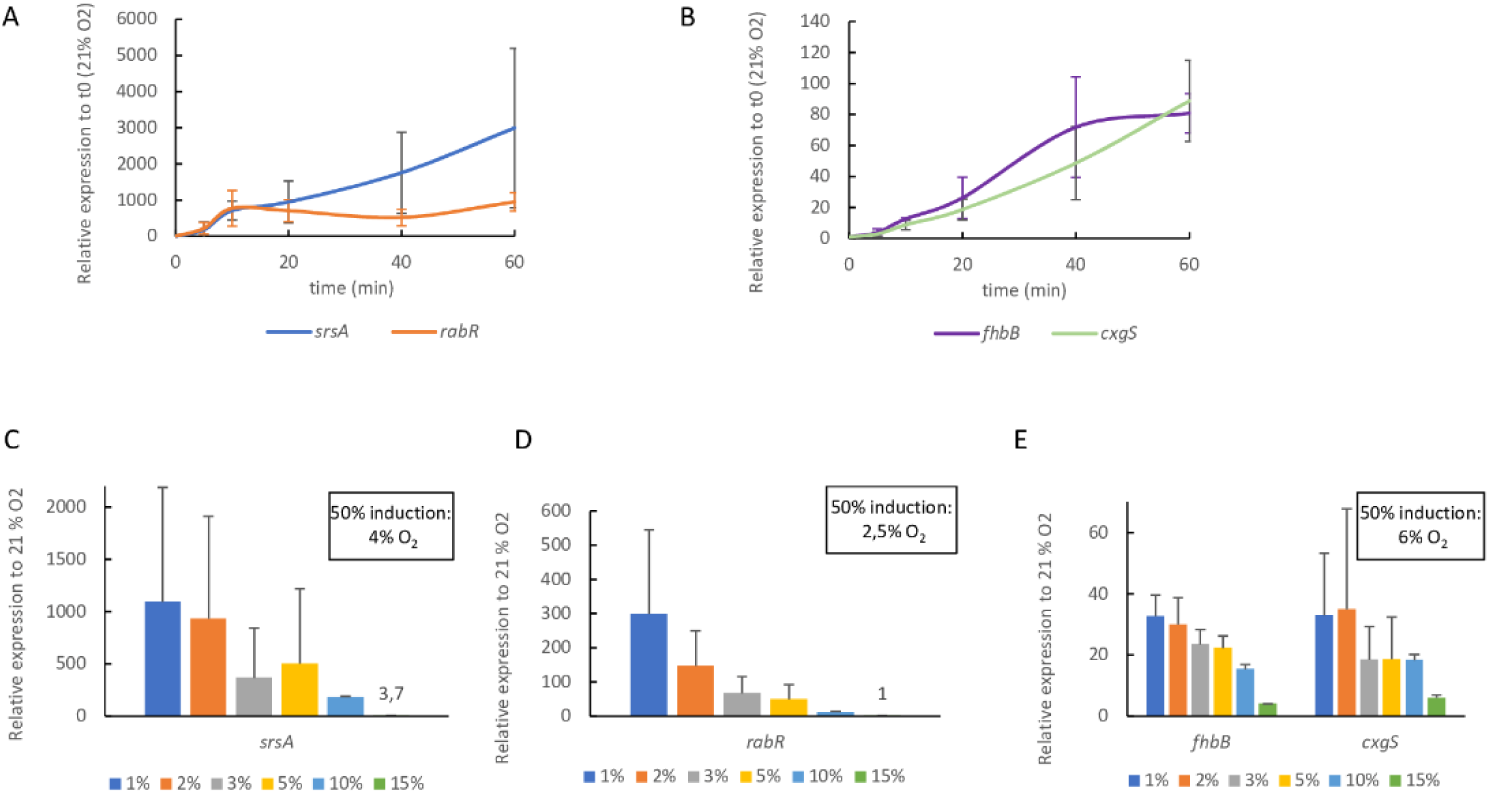
The transcriptomic response to hypoxia relies on multiple and diverse mechanisms. **A and B:** Relative expression of hypoxia marker genes over a short time period (n=4). Cells were placed at 1% O_2_ for 1 h, and collected at the indicated time points. Relative expressions were determined by RT-qPCR using t0 (21% O_2_) as standard. Mean values from 4 trials are indicated with their associated standard deviations. **C-D:** Effect of the oxygen concentrations on hypoxia marker gene expression. Cells were exposed to the indicated concentration of O_2_ for 1 h. Relative expressions were calculated relative to the 21 % O_2_ condition. Mean values from 3 trials with their standard deviations are indicated.

Various inhibitors were tested to probe potential molecular mechanisms that regulate the early inductions of *srsA*, *rabR*, *cxgS* and *fhbB*. As shown in Table S9, DMOG, COCl_2_ and deferoxamine, which inhibit non-heme dioxygenases and demethylases that mimic hypoxic transcriptional effects in other organisms, failed to induce significant expression of these markers, suggesting that they are not controlled by this class of O_2_-sensors. At 1% O_2_, neither DMOG nor COCl_2_ affected gene expression (Table S10). However, DMOG partially lowered *srsA* and *cxgS* expression, reducing their levels to approximately 35% of their normal expression. This suggests that srsA and cxgS expression partially relies on non-heme dioxygenases. The potential involvement of mitochondria was examined using Antimycin A (AA), an inhibitor of the mitochondrial cytochrome C oxidase subunit III (Table 1). At 21% O_2_, AA did not induce gene expression, but it prevented *cxgS* induction under hypoxia. This finding implies a potential role of mitochondrial activity in the expression of the hypoxic cytochrome C subunit. Finally, the involvement of protein synthesis was tested using cycloheximide. This drug had no effect on gene expression at 21% O_2_, but did clearly block *srsA* induction under hypoxia reducing its expression to 1% of the control level. Among the 4 tested genes, only *srsA* induction is dependent on *de novo* protein synthesis.

**Table 1:**
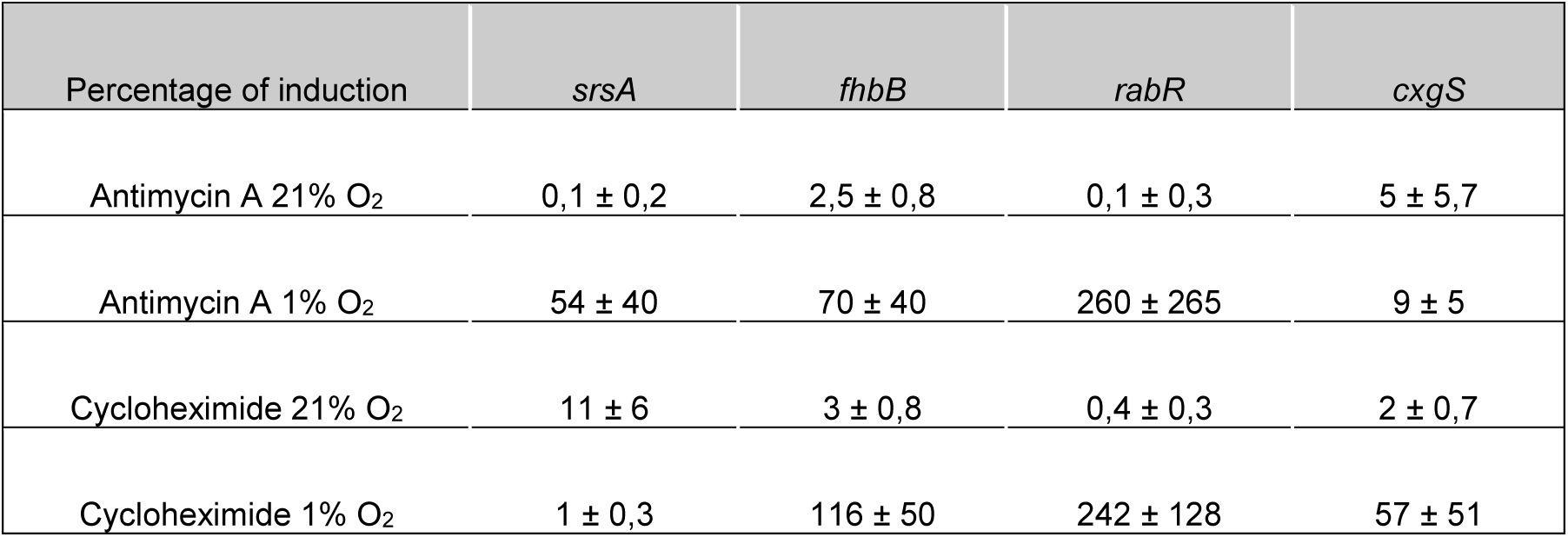
Effects of antimycin A and cycloheximide on genes expressed at 1% and 21% O_2_. Cells were inoculated with the different chemicals for 1 h at either 1% or 21% O_2_. For each gene, relative expression was determined using the 2^-ΔΔCt^ method, using *rnlA* as a reference gene. For cells placed at 21% O_2_, the relative expression was calculated using the condition without chemical treatment as the reference. For cells placed at 1% O_2_, the relative expression was determined using the same chemical condition at 21% O_2_ as the reference. Finally, the percentage of induction at both 21% and 1% O_2_ was calculated using the hypoxic expression level without the drug as the 100% expression. Mean values from 3 trials with their associated standard deviations are indicated.

| Percentage of induction | *srsA* | *fhbB* | *rabR* | *cxgS* |
| --- | --- | --- | --- | --- |
| Antimycin A 21% O<sub>2</sub> | 0,1 ± 0,2 | 2,5 ± 0,8 | 0,1 ± 0,3 | 5 ± 5,7 |
| Antimycin A 1% O<sub>2</sub> | 54 ± 40 | 70 ± 40 | 260 ± 265 | 9 ± 5 |
| Cycloheximide 21% O<sub>2</sub> | 11 ± 6 | 3 ± 0,8 | 0,4 ± 0,3 | 2 ± 0,7 |
| Cycloheximide 1% O<sub>2</sub> | 1 ± 0,3 | 116 ± 50 | 242 ± 128 | 57 ± 51 |

Finally, we investigated the dependence of marker gene transcript levels on O_2_ concentration. As shown in Fig. 9C-E, the expression levels of all the tested genes increased as the O_2_ concentration decreased, with maximum induction apparently occurring at 1-2% O_2_. *srsA*, *fhbB* and *cxgS* showed a significant induction at even at 15% O_2_, while *rabR* induction started at or below 10%. Half maximum induction was observed at 2.5-4% O₂ for *rabR* and *srsA* (Fig. C and D), and at 6% O₂ for *fhbB* and *cxgS* (Fig.9E). The data suggest that gene expression is continuously and inversely proportional to O_2_ concentration, rather than exhibiting switch-like dependence on a critical oxygen concentration threshold. Moreover, *srsA* and *rabR* require lower O_2_ levels than *cxgS* and *fhbB* for maximum induction.

Taken together, the results show that the transcriptomic response to hypoxia is driven by various factors, including time after hypoxia and O_2_ concentration, as well as multiple mechanisms such as *de novo* protein synthesis and mitochondrial activity.

## Discussion and Conclusions

This study showed that *Dictyostelium discoideum* vegetative cells present a strong and complex transcriptomic and proteomic response to hypoxia and reoxygenation. 32% of the detected transcripts were regulated ≥2-fold by hypoxia and 40% of the detected proteins were significantly regulated ≥1.2-fold. This proportion of regulated transcripts is notably higher than what has been reported in other eukaryotes. Studies on *Saccharomyces cerevisiae* identified around 800 differentially regulated transcripts out of total 6,200 genes, and human cell lines show approximately 1,700 regulated transcripts out of 24,000 genes in response to hypoxia (55,56). The RNA turnover between time points was estimated at 20%, which can be compared with the 35% that occurs after 6 hours of starvation-induced development (57). This study is focused on mRNA, but TPM analysis also revealed that the non-coding RNA *dutA*, which is an inhibitor of late development (58), was particularly induced between 5 and 24 h of hypoxia, representing more than 1% of the total amount of detected RNA within the cells. Thus, ncRNAs may also play a role in the transcriptomic regulation of the hypoxic response.

By identifying and analyzing a group of marker genes, we observed extremely rapid (≤10 min) responses for *srsA* and *rabR* at the transcriptional level that were also quickly followed at the protein level. These results may suggest an important role for these proteins in responding to hypoxic conditions. Their responses to hypoxia are also graded with increasing induction over the range of 10 to 1% O_2_. The halfway point varied modestly between genes, suggesting multiple control pathways. The rapid expression of *srsA* relies on *de novo* protein synthesis, suggesting that it may be regulated by a HIF-like mechanism, where a protein is constantly synthesized but degraded under normoxic conditions. By contrast, RabR induction is as rapid but does not require *de novo* protein synthesis, pointing toward existence of another, fast-acting, transcription regulation mechanism. Regarding *cxgS* induction, a study in mammals showed that HIF-1 expression is also inhibited by AA and other mitochondrial chain inhibitors (50), so may be related. Considering the potential importance of non-heme dioxygenases as O_2_-sensors, we tested their potential involvement using the pan-specific inhibitors cobalt (COCl_2_), deferoxamine and DMOG. However, these revealed no effect on early marker transcript expression, though roles in the chronic phase are not excluded. Of specific interest, the Skp1 prolyl hydroxylase PhyA that regulates O_2_-regulation of culmination was down regulated 2-3-fold during both phases, supporting absence of its involvement in the response studies here.

The RT-qPCR results for the data summarized above showed high standard deviation values, indicating significant variability, even though the same induction pattern was consistently observed. The source of this variability was investigated by comparing the variability of the Ct at t0 and 1 h after hypoxia (Fig. S5). The results revealed that the major variability occurred at t0. A similar pattern was observed with RNA-seq data when comparing the TPM at t0 and 1 h post-hypoxia. These data suggest that, despite controlling the cell culture conditions (density, temperature, agitation), and the harvesting procedures, minor experimental fluctuations can lead to large expression changes, especially for sensitive genes like *srsA* or *rabR*. This might reflect biological adaptation to slightest changes that can occur in a natural environment.

The longer term hypoxic response strongly depends on time after hypoxia and can be summarized as occurring in two major phases for both the transcriptome and the proteome: acute (1-5 h) and chronic (24 h). We found that the proteomic data were only moderately positively correlated with the transcriptomic data during chronic hypoxia, while acute hypoxia showed poor correlation. Indeed, we observed that the early transcriptomic adaptation (1 h) was associated with an induction of transcripts involved in ribosome biogenesis, which was not reflected at the protein level. Possible explanations are that ribosome-related proteins experience increased turnover necessitating increased synthesis during hypoxia, or that mRNAs are sequestered in stress granules, which are present in many eukaryotes (59). The observed increase in total mRNA upon hypoxia fits better with the second hypothesis. The role of ribosomes under stressful conditions remains unclear in most eukaryotes and might involve complex regulation. In yeast, for example, ribosome biogenesis is halted, and global translation is inhibited during stress (60). In mammalian cells, a recent study revealed that rRNAs can be produced and stored within the nucleus before re-entering the ribosome biosynthesis pathway once the stress is resolved (61). In contrast, histone proteins drop during chronic hypoxia, while their mRNA levels remained stable. 90% of the histones are typically synthesized during the S phase (62). In our case, hypoxia seems to arrest the cell cycle, which may offer a potential explanation for this down-regulation. Another hypothesis is that histone degradation could induce heterochromatin reorganization, which may explain the large amount of regulated gene expression observed during chronic hypoxia.

We showed that the early proteomic regulation at 5 hours was characterized by an increase in O_2_-binding and redox-related proteins and lipid metabolism. Previous studies have shown that in mammalian cells, DesA is an O_2_-dependent enzyme expressed under hypoxic conditions, leading to the accumulation of bioactive dihydroceramide substrates suited for low O_2_ environments (63). The hypoxia-induced activation of DesA in mammalian cells was interpreted as a compensatory mechanism to mitigate the effects of hypoxia, and this interpretation might also apply to this and the other up-regulated DEPs in *Dictyostelium*. Notably, four lipid-processing-related proteins were found to potentially participate in sterol biosynthesis. Some of the enzymes involved in sterol biosynthesis are O_2_-dependent, and their products regulate the membrane-bound sterol regulatory element-binding protein (SREBP) transcription factor, which in turn controls the majority of hypoxia response genes in certain fungi and yeast (64). It remains unclear whether these observed responses suggest the existence of a similar hypoxia response mechanism in *Dictyostelium* or if they represent a compensatory adaptation for sterol biosynthesis.

The later transcriptomic and proteomic responses are linked to an activation of the cAMP pathway that is reminiscent of early starvation induced development and may enable a collective escape from stressful conditions by promoting migration to a more oxygen-rich environment. Hypoxia could also act as a trigger for *Dictyostelium* sexual reproduction, which relies on cAMP signals and occurs in submerged environments, typically characterized by low oxygen levels. Moreover, *tmcB*, which plays a key role in macrocyst formation and regulates the cAMP pathway during sexual reproduction (65), was slowly induced after 24 hours of hypoxia (Table S1B). In addition, *aco*, which produces ethylene essential for sexual reproduction (66), was strongly induced after 24 h and during reoxygenation. However, this dioxygenase has also been shown to be crucial for *Dictyostelium* culmination (67), so is not specific to macrocyst formation. Although some of the corresponding proteins were detected in our proteome screen, none met our criteria for quantification owing presumably to low expression levels. Alternatively, the increased levels of proteins involved in Ca^2+^-response, adhesion and motility might support the previously described microphase separation of *Dictyostelium* under hypoxia (68).

This study also emphasizes a reduction in cell division after 5 hours of hypoxia, followed by a recovery period of more than 5 hours of reoxygenation, which aligns with previous observations made in *Dictyostelium* (68). This reduction was associated with inhibition of the expression of cell cycle transcripts and proteins specifically during chronic hypoxia, and include transcripts associated with S phase and metaphase-to-anaphase transitions. Arrest also occurs in mammals, where HIF induces the cyclin-dependent kinases inhibitors p21 and p27 (69–71). Other studies also described a HIF-independent mechanism, where PHD1 can modify Cep192 involved in centrosome duplication, leading to its proteasomal degradation (72,73).

Interestingly, most transcripts and proteins regulated during reoxygenation showed continuity with their expression patterns observed during chronic hypoxia. This suggests that reoxygenation does not activate specific metabolic pathways on its own. This contrasts with what is observed in mammals, where reoxygenation induces a rapid increase in antioxidant and stress-related proteins to face the burst of O_2_ (74). Moreover, these data suggest that most transcript and protein levels require more than 5 h of reoxygenation to return to their basal state after exposure to chronic hypoxia. This could represent a preventive strategy to cope with another episode of hypoxia.

Finally, we compared our 1 h transcriptomic data with data from *Dictyostelium* cells exposed to different bacteria or folate (as proxy) for 4 hours (45). Comparison of differentially expressed transcripts showed that GPCRs involved in bacterial sensing displayed a similar pattern under hypoxic conditions, including the down-regulation of *grlA*, *grlB* and *grlG* (Fig. S6A). The hypoxic response marker genes, *rabR*, *srsA*, *cxgS*, *cxgE*, *fhbA* and *fhbB*, were also induced or repressed with the same pattern in presence of bacteria (Fig. S6B). Oxygen depletion by bacterial respiration might explain the common characteristics of the responses. Interestingly, SrsA was previously described to be involved in the early (1 h) starvation response (75), suggesting a connection with nutritional stress that deserves to be explored in future studies.

In conclusion, these results reveal the complexity of the hypoxic response by highlighting the biological processes activated by hypoxia and investigating the multiple factors that drive this response. This study opens the door to further research on hypoxic regulation in *Dictyostelium discoideum*.

## Data availability

Total transcriptomic data will be available on NCBI and DictyExpress (https://app.dictyexpress.org/). The proteomic data (summarized in Table S6) are deposited in the ProteomeXchange Consortium via the PRIDE (48) partner repository with the dataset identifiers PXD061931, PXD062041, PXD061803, PXD061768, PXD061993.

## Supplementary data

Supplementary Data are available at NAR Online.

## Acknowledgements

We thank A. Kafousi for help in RT-qPCR measurements, and S. Sawai for helpful discussions.

## Author contributions

Conceptualization: C.A., C.M.W., J.H. / Methodology: C.A., C.M.W., J.H., S.J. , E.G.P. / Formal analysis: J.H., E.G.P., O.C.E. / Investigation: J.H., H.v.d.W, C.M.W., C.A./ Writing—original draft preparation: J.H, C.M.W. / Writing—review and editing: J.H., C.M.W., C.A. / Visualization: J.H., C.M.W, E.G.P., C.A. / Funding acquisition: J-P.R., C.M.W.

All authors have read and agreed to the published version of the manuscript.

## Funding

This work was supporting by the International Human Frontier Science Program Organization, Grant Number RGP0051/2021 (to J.-P. Rieu, C.M. West).

## Conflict of interest

The authors declare no conflict of interest.

---

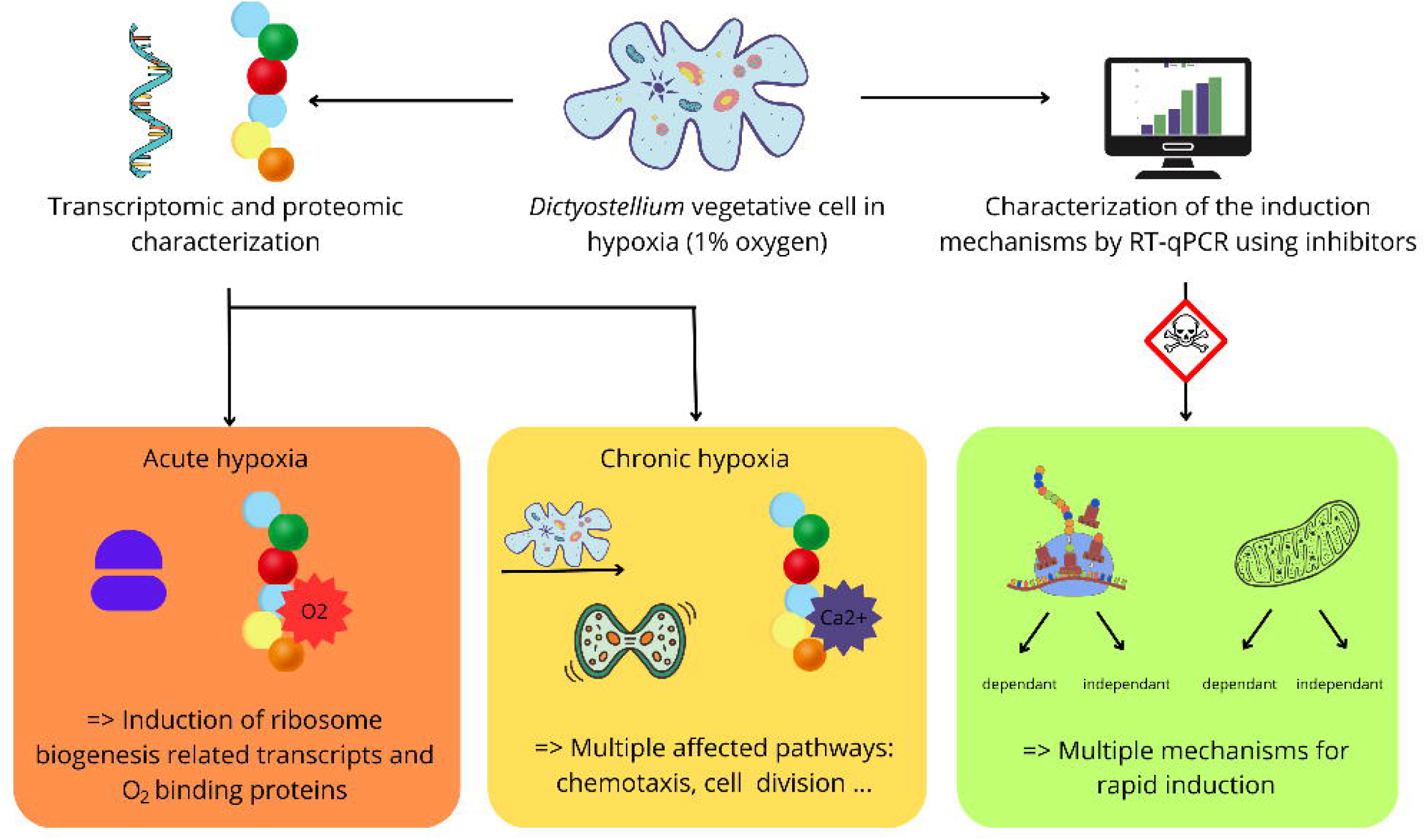

## Notes

### Competing Interest Statement

The authors have declared no competing interest.

